# Hyperexcitable but seizure-free: hippocampal network dysfunction in a Wolfram syndrome model

**DOI:** 10.64898/2026.09.27.754840

**Authors:** Saumel Ahmadi, Juan Gallardo Pinera, Sophie Dokholyan, Andrew Sauerbeck, Hong Nhung Phan, Brianna Carman, Avishek Debnath, Samuel Brunwasser, Gabriel Skinner, Nicholas Rensing, Michael Wong, Cory Cearlock, Susan E. Maloney, Terrance Kummer, Srikanth Singamaneni, John Cirrito, Thomas Foutz, Fumihiko Urano

## Abstract

Wolfram syndrome (WS) is a rare, life-limiting progressive neurodegenerative disorder caused by pathogenic variants in the *WFS1* gene. It is a prototype endoplasmic reticulum (ER) stress disorder, and the mechanisms underlying neurological phenotypes in WS remain incompletely understood. Although WFS1 protein is abundant in the hippocampus, the role of this region in WS remains relatively understudied. Here we characterize hippocampal function across scales in mice carrying a patient orthologous *Wfs1* variant (p.W542*) on both alleles. Male mutants lose excitatory synapses, and dissociated hippocampal neurons are markedly hyperexcitable yet generate no increase in network bursting and no shift in network criticality, indicating that their additional firing is not recruited into coordinated population activity. This finding translates *in vivo* wherein intracranial electroencephalogram (EEG) recordings show no epileptiform activity or seizures in either sex, with no hippocampal astrogliosis, alongside a male-specific reduction in hippocampal low-gamma power. Additionally, EEG demonstrates a sex-specific disruption of sleep, which parallels a sex-divergent synaptic phenotype. Mutant male mice show impaired hippocampus-dependent fear memory and reduced voluntary motor activity. Taken together, we propose that cellular hyperexcitability in the WS hippocampus does not translate into increased network bursting, potentially due to disproportionate loss of excitatory synapses. This dissociation may provide a functional substrate for the neurological features of WS.

## Introduction

Neurological phenotypes emerge from brain circuits, which are in turn built from cellular and intracellular signaling. In channelopathies like *KCNQ2*-related epilepsy, membrane excitability is directly affected, and seizures are a defining feature in patients^1–3^. Where synaptic connectivity is altered instead, the phenotypes can diverge. For example, Alzheimer’s disease (AD) is characterized by hippocampal excitatory synapse loss, with seizures occurring only in a minority of patients^4–7^, while in schizophrenia, excitatory synaptic density is selectively reduced onto parvalbumin interneurons^8^, producing a neuropsychiatric rather than epileptic phenotype. Together this suggests that molecular changes can lead to a variety of different phenotypes depending on the nature of circuit dysfunction. Wolfram syndrome (WS) is a monogenic disorder which offers this opportunity. Wolframin (WFS1) protein is enriched in the hippocampus, WS patients present with neuropsychiatric and cognitive rather than epileptic features, and seizures are uncommon despite progressive neurodegeneration. Yet, synaptic and network-level function have never been examined in WS.

WS is a rare genetic, progressive neurodegenerative disorder, predominantly caused by bi-allelic loss-of-function variants in *WFS1* gene, with an estimated prevalence of 1 in 160,000 to 770,000^9,10^. Classically, it is defined by the association of juvenile-onset diabetes mellitus, progressive optic nerve atrophy, sensorineural hearing loss, diabetes insipidus, sleep-disorder, and progressive neurodegeneration. Death occurs at a median age of 30 years, from brainstem dysfunction and respiratory failure^9,11–13^, with no FDA approved disease-modifying treatments, and management remains supportive^9,14^. Even though neurological involvement is a principal cause of disability and death in WS, neuronal, synaptic and network level function has not been characterized in any model system.

*WFS1* gene encodes wolframin, an endoplasmic reticulum (ER) transmembrane protein that regulates the unfolded protein response, ER calcium homeostasis, and the WFS1 protein is enriched at mitochondria-associated ER membranes^15,16^. Hence, WS is considered as a prototype ER stress disorder. Work in primary neurons has established a coherent subcellular mechanism where loss of wolframin protein compromises ER calcium containment: ER calcium stores are depleted, cytosolic calcium rises, ER-to-mitochondria calcium transfer is impaired, and mitochondrial ATP production falls, producing a bioenergetic deficit that damages neuronal health^17^. WFS1 loss also delays neuronal development through inositol 1,4,5-trisphosphate receptor (IP3R) dysfunction and disturbed mitochondrial dynamics^18^. Because cytosolic calcium is a principal determinant of neuronal firing, the increase in cytosolic calcium in WS neurons should increase neuronal excitability. This prediction has not been tested. Also, it remains unresolved how this well-characterized cellular pathology scales up and affects neuronal networks, ultimately leading to the clinical phenotype. Taken together, translation from loss of WFS1 protein at the cellular level to circuit-level dysfunction has not been directly measured in WS.

This knowledge gap in circuit-level function is important to address because the neuropsychiatric burden of the disease is substantial, and mechanistically unexplained. Early reports described depression, psychosis and other psychiatric co-morbidities in majority of WS patients^19^. Importantly, the neuropsychiatric phenotype is not explained by brain imaging findings, which shows preferential atrophy of brainstem and cerebellum, together with white matter abnormalities ^20^. This distribution accounts for ataxia, dysphagia, and respiratory failure, but does not clearly explain early onset neuropsychiatric symptoms of anxiety disorder, obsessive-compulsive spectrum disorder, mood disorder, and sleep-wake disorder^21^. This phenotype implicates limbic and hippocampal circuits. This mismatch is also seen at the protein expression level, wherein two decades of neuroanatomical characterization show high expression of wolframin in hippocampal-limbic circuits^22,23^, with strong expression in CA1^22,23^, a pattern we also observe in this manuscript. In WS, cerebellum has the most conspicuous atrophy but low wolframin expression, while high expressing hippocampus has not been studied directly at the functional level. The hippocampus and cerebellum, however, are reciprocally connected. On one hand, cerebellar output modulates hippocampal activity via gamma band coordination and interneuron firing^24–26^, thereby exerting a suppressive influence over hippocampal seizures^27,28^. Notably, despite progressive cerebellar degeneration, seizures are uncommon in WS patients^9^. On the other hand, hippocampal activity also drives cerebellar engagement with hippocampal discharges affecting Purkinje cell firing, and hippocampal theta entraining cerebellar theta^29–31^. This cross-talk between hippocampus and cerebellum raises the possibility that hippocampal dysfunction may contribute to cerebellar degeneration seen in WS. This is also supported by studies which show that Purkinje cells may degenerate from disruption to long range circuits without direct insult to cerebellum^32,33^. It does not by itself establish where the primary pathology lies, since expression is not a sufficient determinant of regional vulnerability, but the concentration of wolframin protein in the hippocampus, together with neuropsychiatric phenotypes implicating the hippocampal-limbic circuits, provides a strong rationale for interrogating hippocampal function directly in WS.

The neurological severity varies markedly between WS patients and is only partly explained by genotype^11,34,35^. Truncating (frameshift/nonsense) variants are broadly associated with earlier, more severe disease than in-frame variants with patients carrying two nonsense/frameshift alleles developing diabetes mellitus and optic atrophy significantly earlier than those with one or none^9,11,34^. This is consistent with our use of a homozygous non-sense allele to model the severe end of the spectrum. The mouse model used in this study bears *Wfs1* p.W542* variant on both alleles which is orthologous to human pathogenic variant *WFS1* p.W540* (c.1620G>A; ClinVar VCV000929945.6).

Sex is another possible source of unexplained phenotypic variance which has received little attention at the level of brain-network physiology. Sex differences in WS are, in fact, already apparent clinically with males developing diabetes mellitus significantly earlier than females in the same registry cohort^11^ and male *Wfs1*-knockout mice present with more severe systemic phenotypes than females^36^. However, network-level and electrophysiological phenotypes have not been examined in a sex-stratified fashion.

In this manuscript, we characterize the neuronal, synaptic, and network phenotypes in a patient-relevant model of *WFS1*-Wolfram syndrome, childhood-onset type^9^, using 129Sv/Ev mice homozygous for *Wfs1* p.W542*. Because disease is more severe in males, certain experiments were primarily performed in males. We show that the male mutant hippocampus loses excitatory synapses, and that dissociated hippocampal neurons are markedly hyperexcitable yet generate no increase in network bursting and no shift away from a near-critical state indicating that the additional excitation they produce is not recruited into coordinated population activity. These findings persist *in vivo*, wherein intracranial electroencephalogram (EEG) did not show any epileptiform activity or seizures in either sex, in the absence of hippocampal astrogliosis, which is concordant with clinical picture. In males, EEG additionally revealed a region-specific reduction in hippocampal low-gamma power. These findings indicate that hyperexcitable hippocampal neurons are embedded in a network with reduced excitatory connectivity, such that hyperexcitability at the unit level fails to translate into network-level hyperexcitability, identifying a failure of excitation to propagate as a potential substrate for the non-epileptic neurological and neuropsychiatric features of WS.

## Results

### A homozygous knock-in WS mouse model carrying the patient-relevant *Wfs1* p.W542* reproduces a WFS1-null state without overt diabetes

We generated 129Sv/Ev mice carrying the patient-orthologous *Wfs1* p.W542* nonsense variant by CRISPR-Cas9–mediated homology-directed repair, introducing a single g>a substitution that converts the wild-type TGG codon at position 542 to a premature TGA stop in exon 8 resulting in NM_011716.2(*Wfs1*): c.1626G>A (p.W542*) (Figure 1a, b). Amino acid numbering follows mouse wolframin (UniProt P56695) the position corresponds to the human pathogenic variant p.W540*. In hippocampus, *Wfs1* transcript measured by RT-qPCR with an amplicon spanning exons 7–8, 5′ of the premature termination codon, was reduced to undetectable levels in homozygous mutants relative to wild-type (n=3 per genotype; p=0.02, Welch’s t-test; Figure 1c). The premature stop lies in exon 8 (terminal exon), and results in near-complete transcript loss, indicating that the allele is not merely destabilized but effectively null at the RNA level. Consistent with transcript loss, immunoblotting with an antibody against an N-terminal epitope (residues 2–400, upstream of the truncation) detected full-length wolframin (∼100 kDa) in wild-type but not mutant hippocampal lysates (Figure 1d), confirming absence of both full-length and any N-terminally retained truncated product and establishing the allele as a functional null. Immunofluorescence showed wolframin protein enrichment in hippocampal CA1 in wild-type (WT) tissue that was absent in mutants (Figure 1e, f); cerebellar wolframin was low to undetectable in both genotypes (Figure 1g, h), confirming that the region bearing the disease’s most prominent atrophy is not a site of high wolframin expression.

**Figure 1:**
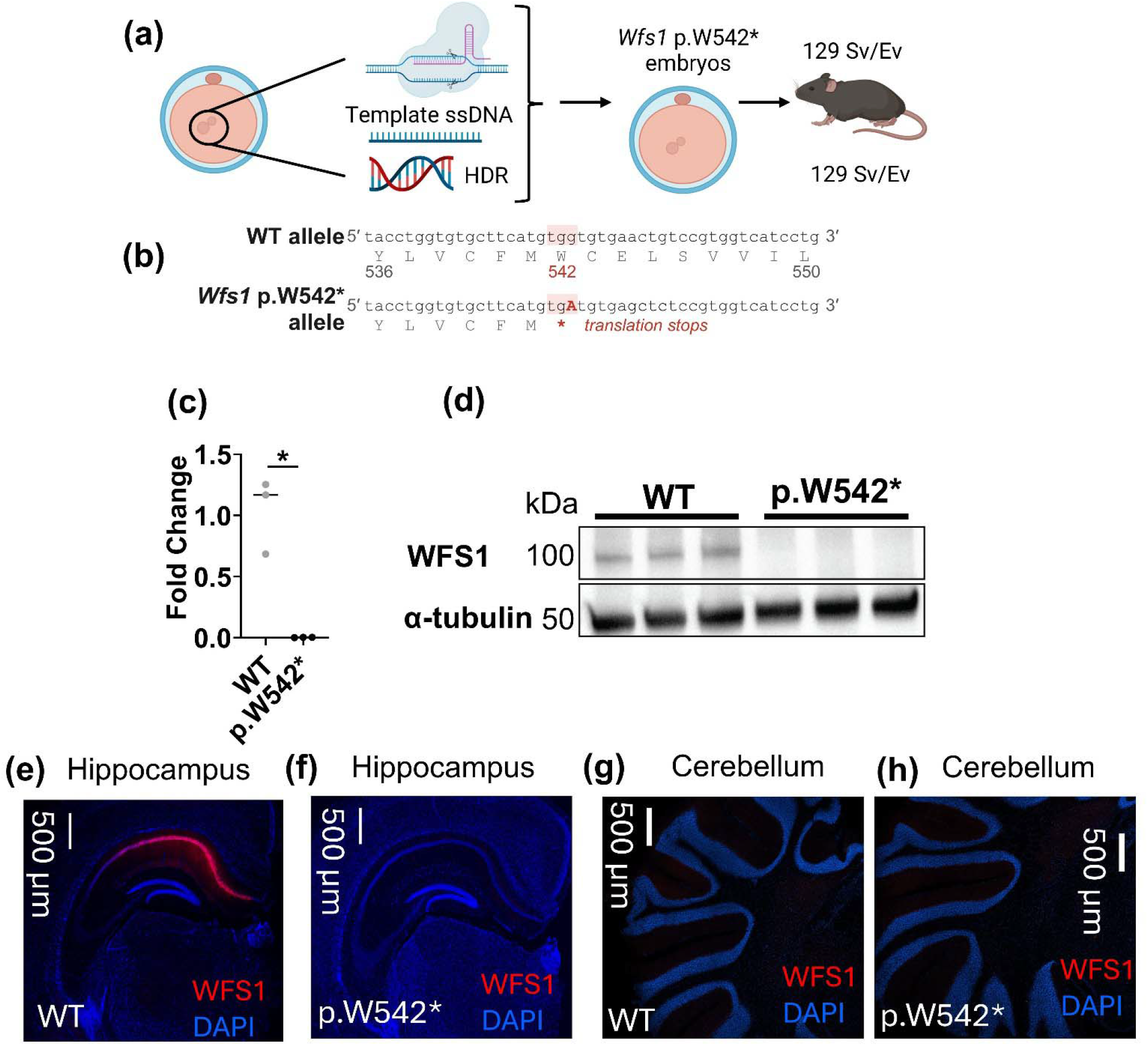
A knock-in Wolfram syndrome mouse model carrying the patient-derived *Wfs1* p.W542* lacks *Wfs1* transcript. **(a)** Generation of the *Wfs1* p.W542* allele. CRISPR-Cas9 ribonucleoprotein and a single-stranded DNA donor template were introduced into 129Sv/Ev zygotes, and the point substitution was made by homology-directed repair (HDR). Founders were backcrossed to 129Sv/Ev and maintained on a congenic 129Sv/Ev background. **(b)** Genomic sequence of the targeted region in the wild-type and p.W542* alleles. The wild-type TGG codon at position 542 (red) is converted to TGA by a single g>a substitution, introducing a premature termination codon in exon 8. Amino acid numbering follows mouse wolframin (UniProt P56695); the equivalent human residue with a known pathogenic variant is *WFS1* W540* (ClinVar VCV000929945.6). **(c)** Wfs1 transcript abundance measured by RT-qPCR in hippocampus from adult Wolfram syndrome mice homozygous for p.W542* variant (p.W542*), expressed as fold change relative to the wild-type mean (ΔΔCt, normalized to GUSB). Amplicon spans exons 7-8, 5′ of the premature termination codon. Each symbol represents one animal; bars indicate group means. n=3 per genotype (*p=0.02, Welch’s t-test). **(d)** Immunoblot for wolframin (∼100 kDa) in hippocampal tissue lysates from three wild-type and three homozygous p.W542* mice, with α-tubulin (∼50 kDa) as a loading control. The antibody used (Abcam ab259362) recognizes an N-terminal epitope within residues 2–400 (exact site propriety to AbCam), which lies upstream of the p.W542 truncation site. **(e–h)** Immunofluorescence for wolframin (red) with DAPI counterstain (blue) in sections from wild-type and homozygous p.W542* mice. **(e, f)** Hippocampus sagittal slices show wolframin immunoreactivity is enriched in CA1 of the hippocampus in wild-type (WT) tissue and is absent in mutants (p.W542*). **(g, h)** Cerebellum sections show wolframin immunoreactivity is low to undetectable in both genotypes. Sections from both genotypes were processed in parallel, imaged with matched acquisition settings, and displayed with identical brightness and contrast.

Homozygous mutants were lighter than WT littermates across 6–20 weeks in both sexes (males −3.65 g, p=1.3×10□□; females −4.07 g, p=2.1×10□¹³; linear regression with age covariate, errors clustered on dam) and shorter in naso-anal length (males −0.38 cm; females −0.39 cm; both p<10) (Supplementary Figure 1a, b, d, e). Notably, fasting blood glucose in 16-week-old males did not differ from wild-type (p>0.99; Supplementary Figure 1c), suggesting that the neurological and network phenotypes described below may emerge in the absence of overt diabetes at the ages studied.

### Hippocampal WFS1 loss in males predominantly reduces excitatory synapse density

To determine whether wolframin loss alters synaptic composition, we applied SEQUIN (Synaptic Evaluation and QUantification by Imaging Nanostructure) to CA1 stratum radiatum (SR) and the dentate gyrus molecular layer (ML) in adult male mice (16 weeks old), as previously described^37^. Similar to previously done SEQUIN^37^, we quantified colocalized synapsin–PSD-95 pairs (less than 260 nm apart) as excitatory synaptic loci. For assessment of inhibitory synapses we co-stained with synapsin (pre-synaptic marker) and gephyrin (post-synaptic marker), however the synapsin–gephyrin separation-distance histogram did not resolve a short-distance peak comparable to that obtained with Synapsin-PSD-95, despite no-primary controls showing no detectable gephyrin puncta. We hypothesize that this could be due to differences in the pool of intra- and extra-synaptic gephyrin densities, wherein previously it has been shown that in hippocampal neurons, substantially more extra-synaptic than synaptic clusters of gephyrin exist, with exchange between compartments occurring with changes in inhibitory plasticity^38^. This is consistent with an inhibitory postsynaptic density that is structurally distinct from its excitatory counterpart^39^, and with heterogeneity across inhibitory synapse classes^40^. We therefore used total gephyrin puncta as a measure of the overall gephyrin scaffold pool rather than of inhibitory synaptic loci. Both sets of experiments were done on n=7 wild-type, n=6 mutant; Figure 2d-g. Excitatory synaptic density was significantly reduced in mutants in both regions (Mann-Whitney test): by 31% in SR (p=0.01) and 28% in ML (p=0.03) (Figure 2d, e). Inhibitory gephyrin puncta were less affected with gephyrin reduced by only 5% in SR (p=0.02) and was not significantly changed in ML (p=0.10) (Figure 2f, g). Additionally, western blotting of male hippocampal lysates showed no significant difference in AMPA or NMDA receptor subunit levels (Supplementary Figure 3) indicating that the reduction in excitatory synaptic loci may not be accompanied by a detectable change in bulk glutamate receptor expression. This data, in male mice, supports a marked loss of excitatory synaptic loci most pronounced in SR, without a comparable change in the total inhibitory gephyrin scaffold.

**Figure 2:**
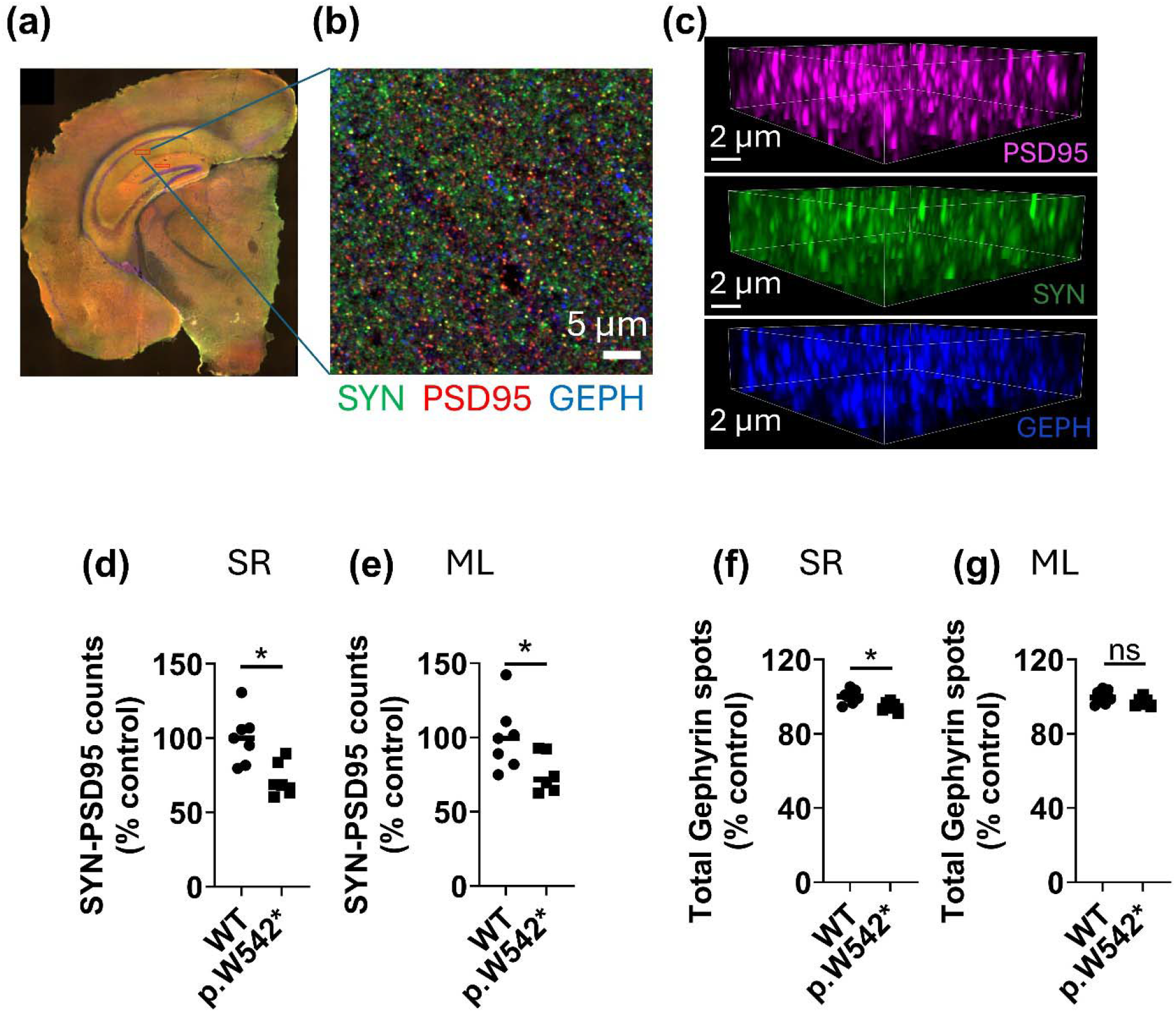
Hippocampal Wfs1 loss in males predominantly reduces excitatory synapse density. **(a)** Representative coronal hemisection from an adult male mouse showing the hippocampus; boxed region indicates the Stratum Radiatum (SR) sampling area imaged at high resolution, along with a second red box in the Molecular Layer (ML). **(b)** Representative single section from SEQUIN (Synaptic Evaluation and QUantification by Imaging Nanostructure) field acquired in SR, immunolabeled for synapsin (SYN, green), PSD-95 (red), and gephyrin (GEPH, blue). **(c)** Three-dimensional volume renderings of the individual channels from a subvolume of the field in (b): PSD-95 (magenta, top), synapsin (green, middle), gephyrin (blue, bottom). **(d, e)** Male mice, SEQUIN quantification of colocalized synapsin–PSD-95 pairs (excitatory synaptic loci) per region of interest. Values are normalized to the wild-type group mean. **(d)** SR of CA1, *p=0.01; difference in median = -31% excitatory synapses per ROI per animal **(e)** ML of the dentate gyrus, *p=0.03; difference in median = - 28% **(f, g)** Male mice, total gephyrin puncta per region of interest in SR and ML. Each symbol represents one animal with each data point representing a mean across 14 ROIs (region of interest) per animal for SR, and 7 ROIs per animal for ML); horizontal bars indicate group medians. Values are normalized to the wild-type group mean. n=7 wild-type (WT) male mice and n=6 p.W542* male mice. Statistical comparisons by Mann-Whitney-test: (f) SR, *p=0.02; difference in median = -5% (g) ML, ns: p=0.10; difference in median = -2.5%.

We next asked if the same synaptic phenotype was seen in female hippocampi (n = 4 WT and n = 5 mutants). We found that co-localized synapsin-PSD-95 loci did not significantly differ in WT vs p.W542* mutants, both in SR (p=0.56) and ML (p=0.73) as shown in Supplementary Figure 4a, b (Mann-Whitney test). However, total gephyrin puncta were reduced in p.W542* mutants in both SR (-9%, p=0.016) and ML (-9%, p=0.016) using Mann-Whitney test. Taken together, these data indicate that the synaptic consequences of loss of wolframin may differ by sex, with excitatory loci predominantly affected in males and inhibitory gephyrin scaffold predominantly affected in females. Interestingly, secondary analyses of intracranial hippocampal EEG data showed that the aperiodic exponent shifted towards inhibition-dominance in male mutants and excitation-dominance in female mutants (Supplementary Figure 5), similar to the observed synaptic phenotypes.

### Hippocampal networks are hyperexcitable but do not increase network bursting

To test how altered synaptic composition translates into emergent network activity, we recorded spontaneous activity from postnatal day-1 (P1) dissociated male hippocampal cultures on multielectrode arrays (MEA) at days *in vitro* (DIV14, n=4 cultures per genotype, each from a separate animal; Figure 3a–d). Mutant cultures were hyperexcitable with weighted mean firing rate per active electrode being elevated nearly threefold (1.96 vs 5.7 Hz; p=0.028; Figure 3e), an increase which was primarily driven by regular-spiking units (putative excitatory neurons; 4.4 vs 9.9 Hz/cell; p=0.028; Figure 3g), with fast-spiking units (putative inhibitory neurons) trending higher but not significantly changed (2.4 vs 5.6 Hz/cell; p=0.68; Figure 3h). Despite this increase in firing, network-level bursting did not increase, and network burst rate was not statistically significant between genotypes (21 vs 29 bursts/min; p=0.88; Figure 3f). We further quantified the network’s operating regime by estimating the branching parameter using the multistep regression (MR) estimator, which is robust to subsampling and to the rate differences between genotypes^41–43^. Across all bin widths, recordings from both genotypes yielded a branching parameter below unity (m<1), placing the networks in a subcritical, near-critical regime with no genotype difference in the distance from criticality (Figure 3i). The conventional single-step branching-ratio estimator was highly bin-width dependent, did not resolve a genotype difference and is reported in Supplementary Figure 2. Taken together, the combination of elevated firing with unchanged network bursting and unchanged criticality defines a state in which hippocampal neurons lacking wolframin are individually hyperexcitable yet their additional firing is not recruited into coordinated, population-level activity.

**Figure 3:**
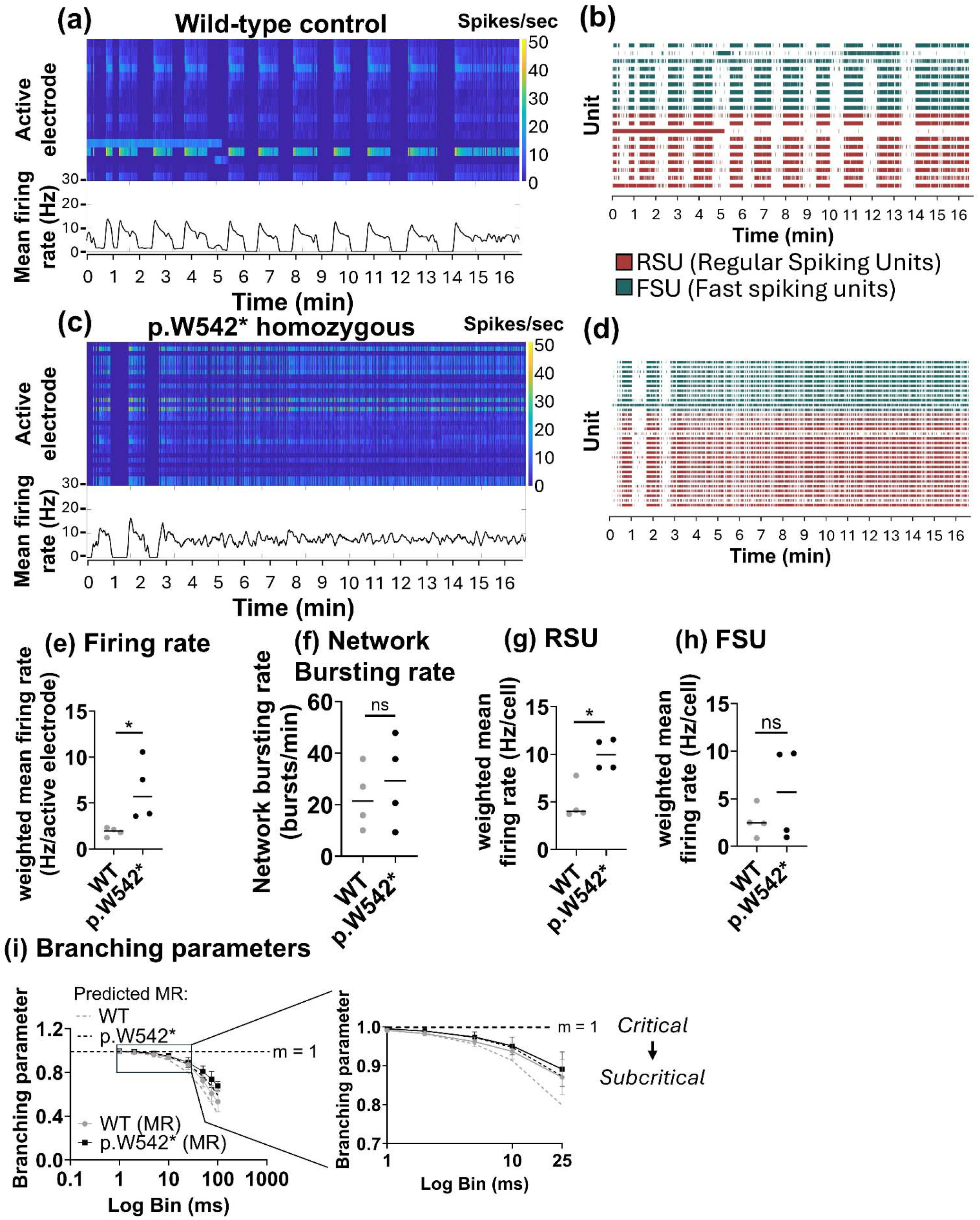
*In vitro* multielectrode array recordings of WS male mouse hippocampal cultures exhibit neuronal hyperexcitability without an increase in network bursting. (**a, c**) Representative multielectrode array (MEA) recordings from post-natal day 1 dissociated hippocampal cultures of a wild-type control (a) and a *Wfs1* p.W542* homozygous (c) male mouse. Recordings were performed at days in vitro (DIV) 14. Top panel shows a raster plot representing firing rate across active electrodes over the recording session (color scale represents spikes/second). Bottom panel shows array-wide mean firing rate (MFR) for the same recording. **(b, d)** Spike raster plots for the recordings in (a) and (c), with sorted units classified as regular-spiking units (RSU, red; putative excitatory neurons) or fast-spiking units (FSU, teal; putative inhibitory neurons). **(e–h)** Group data, n = 4 independent cultures per genotype, each derived from a separate animal, and horizontal bars represent mean values. **(e)** Weighted mean firing rate per active electrode (n=4, WT mean of 1.96 vs p.W542* mean of 5.7, *p = 0.028, Mann-Whitney two-tailed test). **(f)** Network burst rate representing array-wide bursts per minute. Mean for WT 21 vs p.W542* 29 bursts/min; ns: p=0.88, Mann-Whitney two-tailed test **(g)** Weighted mean firing rate per cell for RSU (n=4, mean for WT 4.4 vs p.W542* 9.9 Hz/cell; *p=0.028, Mann-Whitney two-tailed test). **(h)** Weighted mean firing rate per cell for FSU (n=4, mean for WT 2.4 vs p.W542* 5.6 Hz/cell; ns: p=0.68, Mann-Whitney two-tailed test). **(i)** Branching parameter as a function of analysis bin width (1–100 ms), estimated by multistep regression (MR). Data points represent mean +/- standard deviation (SD) with dashed lines representing predicted MR for each genotype fitted for autocorrelation time. The decline of values for branching parameter m is expected for a network with a single intrinsic timescale. Inset to the right shows expansion of the 1-25 ms range. All recordings across both genotypes had m < 1 at every bin width, supporting that both genotypes operated in a subcritical, near-critical regime.

### *In vivo*, WS mice do not generate seizures, and male mice show a low-gamma deficit

To test whether the *in vitro* MEA phenotype has an *in vivo* correlate, we recorded intracranial hippocampal EEG with concurrent nuchal EMG, in 17-19 week old mice, for three consecutive days, across light–dark cycle in both sexes (Figure 4a). Continuous review of these recordings revealed no electrographic seizures or interictal epileptiform discharges in mutants of either sex, concordant with the clinical observation that seizures are uncommon in WS and consistent with a network that is hyperexcitable but sub-critical rather than epileptogenic. Consistent with the absence of epileptiform activity, hippocampal astrocyte reactivity was also unchanged. Glial fibrillary acidic protein (GFAP) immunoreactivity in CA1 stratum radiatum did not differ between genotypes (Supplementary Figure 6). Because reactive astrogliosis is a defining feature of hippocampal sclerosis and correlates with increasing seizure burden^44–46^, its absence likely indicates absence of ongoing epileptogenesis.

**Figure 4:**
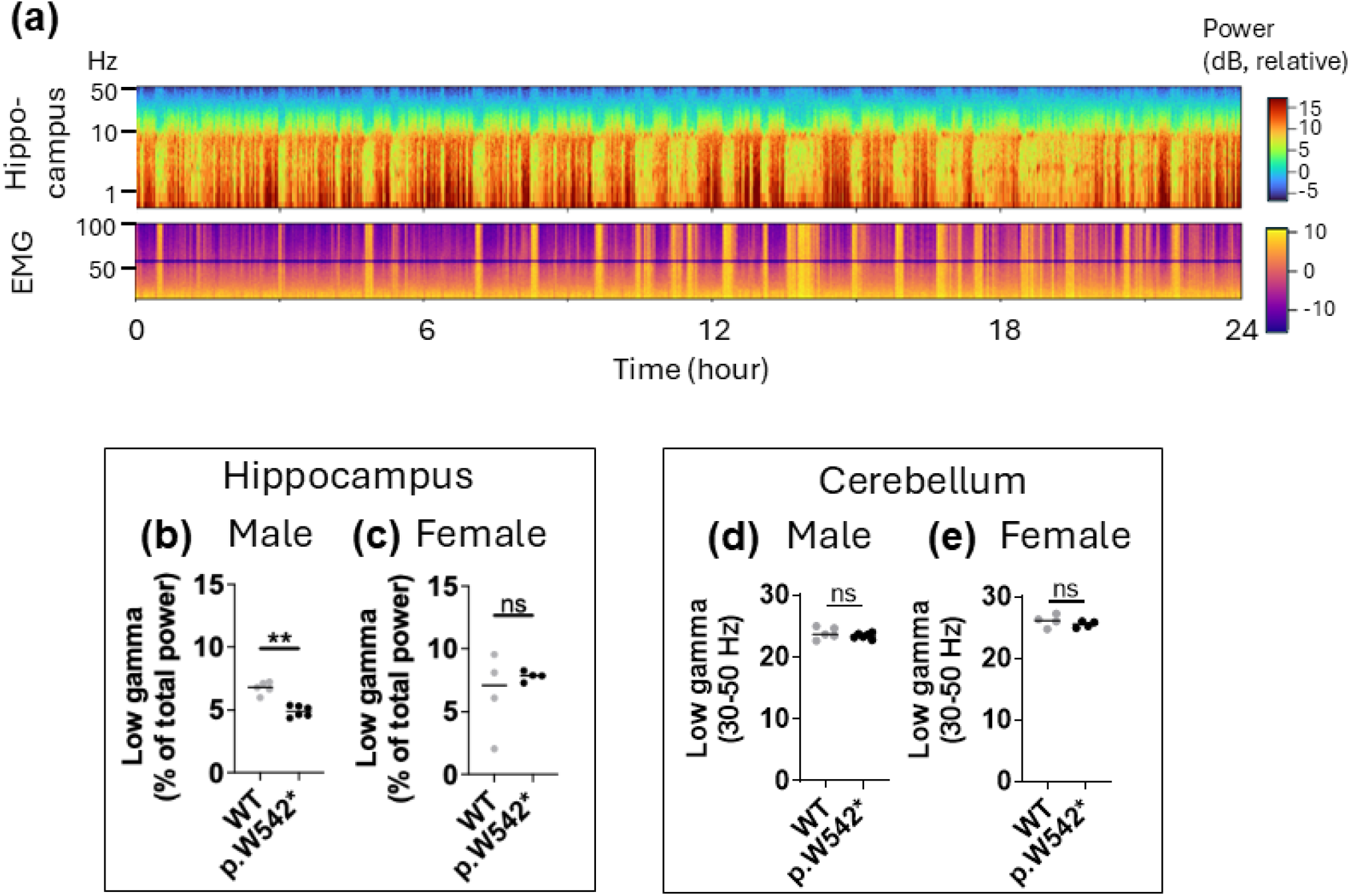
*In vivo* intracranial hippocampal EEG recordings reveal a low gamma deficit in males with absence of epileptiform activity in both males and females. (**a**) Representative 24-hour hippocampal EEG and nuchal EMG spectrograms from a single Wild-type (WT) male animal across one light–dark cycle. Color scale represents relative power in decibels referenced to the median power of the recording; y-axes show frequency (Hz). Recordings were screened for epileptiform activity as described in Methods. (b, c) Hippocampal low-gamma relative power (30–50 Hz; expressed as percentage of total 0.5-90 Hz power) recorded from hippocampus in male mice (b; WT n=5, p.W542* n=6; Mann-Whitney test, **p=0.004, Difference in median of -1.9 percentage points or -29%) and female mice (c; WT n=4, p.W542* n=4; Mann-Whitney test, ns p=0.8, Difference in median of 0.7). (d, e) Similarly, relative low-gamma power recorded from cerebellar depth electrodes in the same male (d) and female (e) animals, with no significant differences in low-gamma power in both males and females.

Spectral analysis revealed that hippocampal low-gamma power (30–50 Hz; expressed as percentage of total 0.5-90 Hz power) was reduced in male mutants (median difference −1.9 percentage points, −29% relative reduction; WT n=5, mutant n=6; p=0.004; Figure 4b) but not in females (+0.8 percentage points, n=4 per genotype; p=0.8; Figure 4c), and the genotype effect differed between sexes (genotype x sex interaction, permutation p=0.027). Because relative power is normalized to the whole spectrum, changes in the aperiodic exponent may be reflected in low-gamma band power (Supplementary Figure 5), rather than an isolated change in gamma oscillatory activity. The deficit was also regionally selective, with cerebellar low-gamma power, recorded from depth electrodes in the same animals, being unchanged in both sexes (Figure 4d, e). The low-gamma reduction is therefore both male-specific and potentially hippocampus-specific. Thus, male mutants exhibit a hippocampal reduction in relative to low-gamma power in the context of altered aperiodic spectral structure. Because gamma-band activity depends on coordinated fast interactions across a connected neuronal population^47^, a selective loss of hippocampal low-gamma is consistent with reduced excitatory synaptic density observed in SR.

### Sleep architecture is disrupted in a sex-specific manner

From the same intracranial EEG recordings, sleep was scored as wake, NREM, and REM (Figure 5a–c) using Somnotate^48^. Sleep disruption differed by sex. Male mutants showed increased REM-to-wake transitions (median difference +16; p=0.02; Figure 5d), unchanged in females (p=0.6; Figure 5e). Female mutants instead showed altered NREM structure with more NREM bouts (p=0.02; Figure 5g) of shorter mean duration (−32 s; p=0.02; Figure 5i), with male NREM bout number and duration unchanged (p=0.3, p=0.93; Figure 5f, h). Sleep is thus disrupted in both sexes but through distinct architectural signatures, wherein, REM-exit occurs more often in males and NREM fragmentation in females. Given that hippocampal gamma activity is coordinated in a state-dependent manner with REM sleep^49,50^, the co-occurrence in males of a hippocampal low-gamma deficit and REM-exit instability may reflect a shared circuit substrate, however further studies would need to be done to determine that.

**Figure 5:**
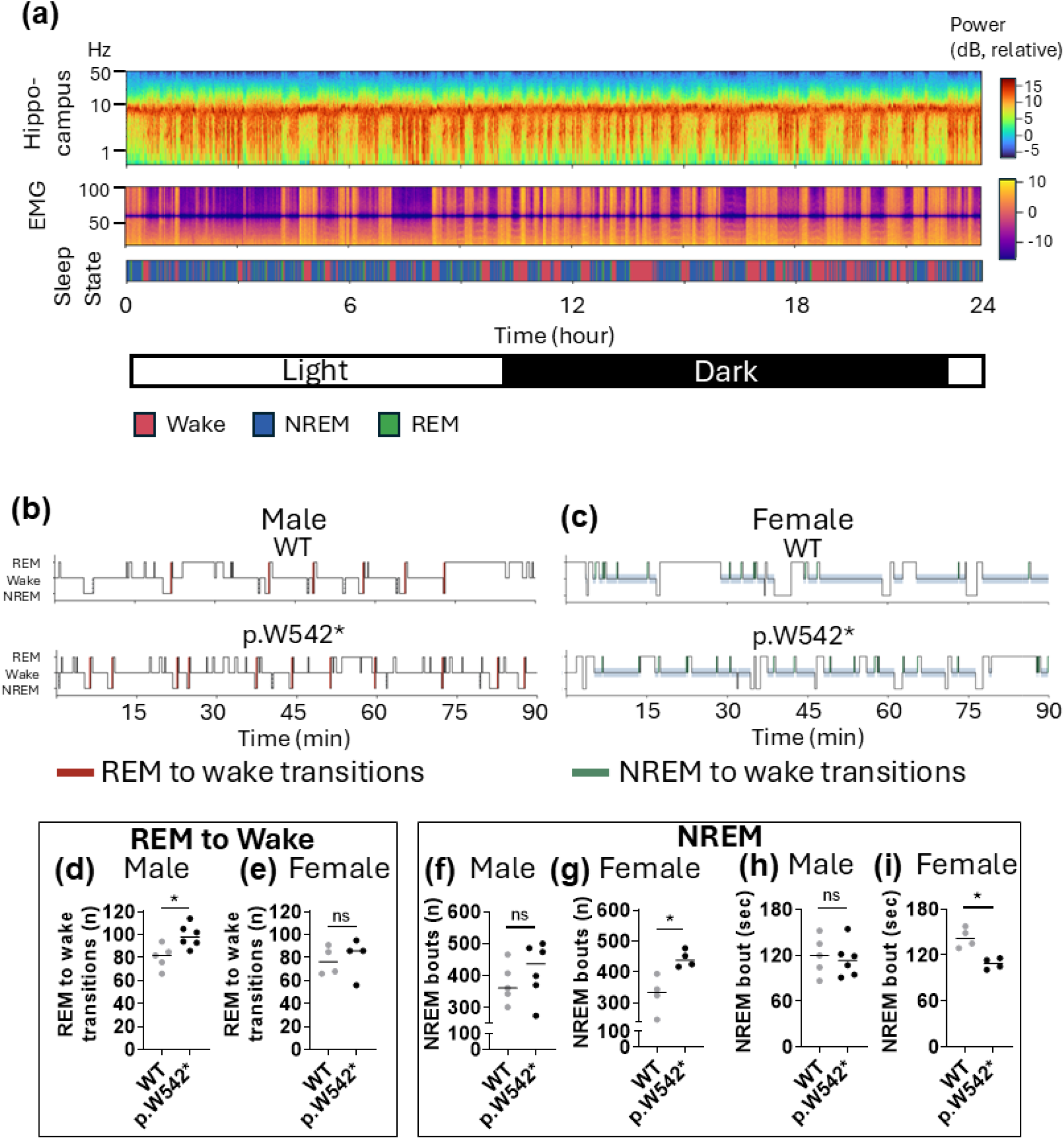
Hippocampal EEG shows altered sleep architecture in Wolfram syndrome mice. **(a)** Representative 24-hour recording from a single Wild-type male mouse. Top, spectrogram of the hippocampal EEG; middle, spectrogram of the nuchal EMG; bottom, corresponding hypnogram scored as REM, wake, and NREM. Frequency axes in Hz; color scale indicates spectral power in dB. The light-dark bar alternating light/dark exposure every 10 hours. **(b, c)** Representative 90 min hypnogram excerpts for (b) male and (c) female wild-type (top) and homozygous p.W542* (bottom) mice. Red marks denote REM-to-wake transitions; green marks denote NREM to awake transitions. **(d–i)** Sleep architecture quantified over full recording per animal. (d, e) Number of REM-to-wake transitions in males (d; WT n=5, p.W542* n=6; Mann-Whitney test, *p=0.02, Difference in median of 16) and females (e; WT n=4, p.W542* n=4; Mann-Whitney test, ns p=0.6, Difference in median of 9.5). **(f, g)** Number of NREM bouts in males (f; WT n=5, p.W542* n=6; Mann-Whitney test, ns p=0.3, Difference in median of 75) and females (g; WT n=4, p.W542* n=4; Mann-Whitney test, * p=0.02, Difference in median of 104). **(h, i)** Mean NREM bout duration in males (h; WT n=5, p.W542* n=6; Mann-Whitney test, ns p=0.93, Difference in median of -6.8) and females (i; WT n=4, p.W542* n=4; Mann-Whitney test, * p=0.02, Difference in median of -32).

### Male mutants show impaired fear memory with intact acquisition, and a selective voluntary motor deficit

Male mutants acquired conditioned fear normally on day 1 (p=0.212, η^2^_p_=.067; Figure 6a, b) but froze significantly less than wild-type littermates to the conditioning context on day 2 (p=0.015, η^2^_p_=0.233; Figure 6c) and to the previously paired tone on day 3 (p=0.010, η^2^_p_=0.255; Figure 6d), indicating deficits in both contextual and cued fear memory in the presence of intact acquisition. Next, we distinguished voluntary from forced motor behavior. Because the activity measures were non-normally distributed, analyses were performed on log□□-transformed scale. In voluntary wheel running, mutants achieved less distance (p=0.0003, η^2^ = 0.452), lower maximum speed (p=0.0002, η^2^ =0.474), and fewer total rotations (p=0.0003, η^2^ =0.456) than wild-type across normal and complex wheel segments (Figure 7a–c). This suggests a robust reduction in self-initiated activity. In contrast, on the accelerating rotarod which is a forced task unadjusted performance p.W542* mutants performed indistinguishably from WT on latency to fall (p=0.44, η²p=0.016), distance travelled (p=0.57, η²p=0.009), and speed achieved (p=0.96, η²p=0.001) (Figure 7d–f), indicating intact motor coordination and balance. It was only after adjustment for body weight as a covariate that we saw reductions in latency to fall (p=0.044, η²p=0.108) and distance travelled (p=0.022, η²p=0.137), with speed unchanged (p=0.19, η²p=0.048) (Figure 7g–i) highlighting the fact that mutants and WT differ substantially in body weight. However, this adjustment extrapolates beyond the observed weight range in each group and hence these covariate-adjusted values should be interpreted with caution. Taken together, these findings suggest that the motor phenotype is characterized primarily by reduced voluntary, self-initiated activity, with relatively preserved performance on a forced motor task.

**Figure 6:**
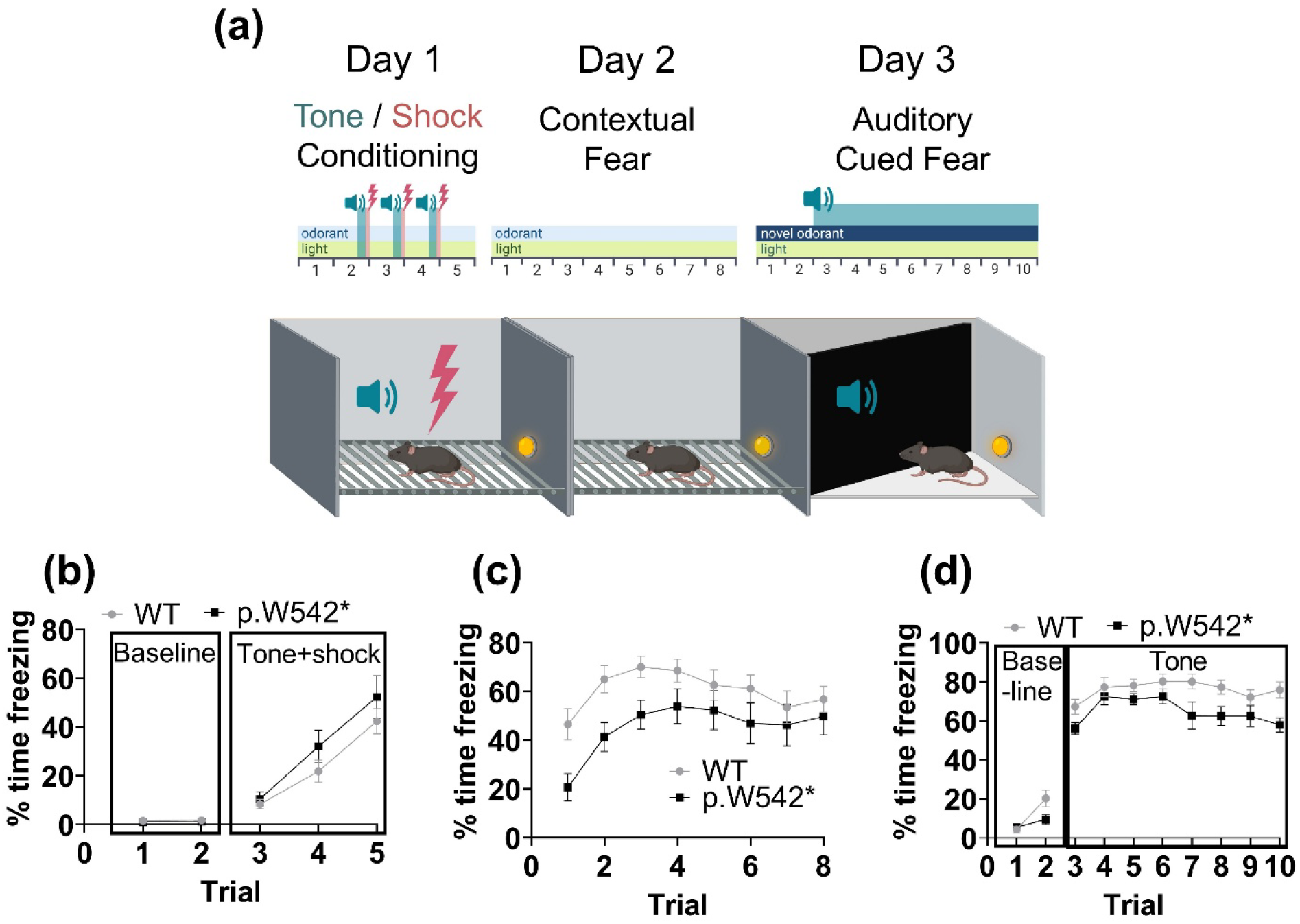
WS male mice show impaired contextual fear retrieval despite intact acquisition. **(a)** Schematic and cartoon of the fear conditioning assay. **(b)** Day 1 freezing levels demonstrate no differences between WT and p.W542* male littermates in their response to a mild foot shock during conditioning (*F*(1,23)=1.647, p=0.212, η^2^_p_=0.067). **(c)** On day 2, male p.W542* mice froze significantly less than WT littermates in response to the context of the prior foot shock presentation (*F*(1,23)=6.985, p=0.015, η^2^_p_=0.233), indicating a contextual fear memory deficit. **(d)** On day 3, male p.W542* mice again froze significantly less than WT littermates in response to the cue tone previously paired with the foot shock presentation (*F*(1,23)=7.878, p=0.010, η^2^_p_=0.255), demonstrating a cued fear memory deficit.

**Figure 7.**
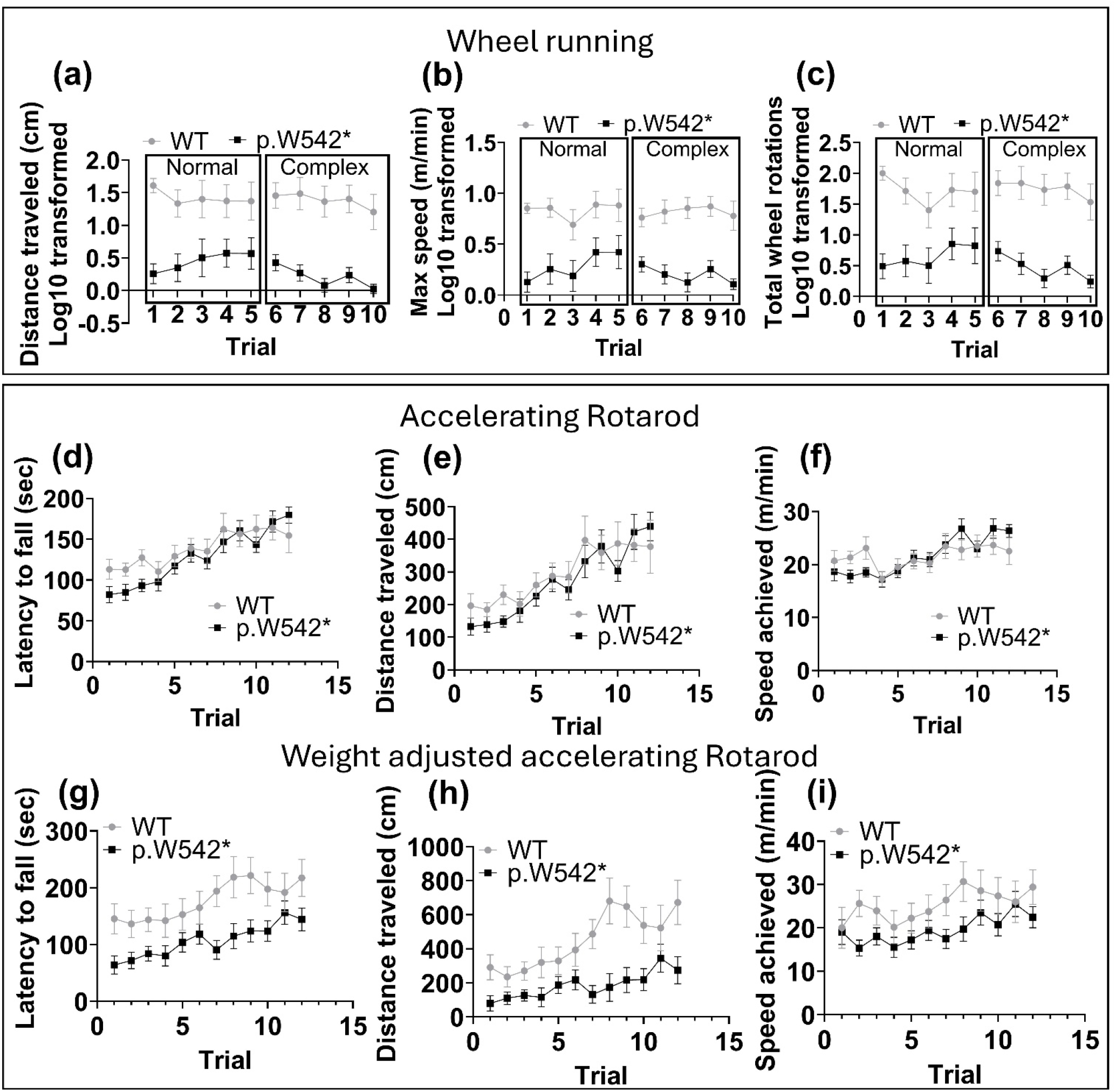
WS male mice show a selective deficit in voluntary motor behavior. **(a-c)** During activity wheel running, p.W542* mutant mice were unable to achieve comparable (a) distance travelled (F(1,22)=10.774, p=0.003, η^2^ (F(1,22)=10.901, p=0.003, η^2^ (F(1,22)=10.936, p=0.003, η^2^ =0.329; F(1,22)=18.111, p=0.0003, η^2^ =0.331; F(1,22)=19.802, p=0.0002, η^2^ =0.332; F(1,22)=18.429, p=0.0003, η^2^ = 0.452) (b) max speed =0.474) (c) total wheel rotations =0.456 to WT littermates across both the normal and complex wheel segments. **(d-i)** On the accelerating rotarod which is a forced task requiring motor coordination and balance, there was no difference between the WT and p.W542* homozygous mice (d) latency to fall (*F*(1,37)=0.609, p=0.440, η^2^_p_=0.016) and (e) total distance traveled (*F*(1,37)=0.321, p=0.574, η^2^_p_=0.009), (f) speed achieved (*F*(1,37)=0.002, p=0.964, η^2^_p_=0.001). **(g-i)** However, with body weight as a covariate, and adjusting for it, leads to extrapolation of values, and shows a reduction in (g) latency to fall (*F*(1,36)=4.356, p=.044, η^2^_p_=0.108) and (h) total distance traveled (*F*(1,36=5.701, p=0.022, η^2^_p_=0.137), yet unchanged in (i) speed achieved (*F*(1,36)=1.822, p=.186, η^2^_p_=0.048).

**Figure 8.**
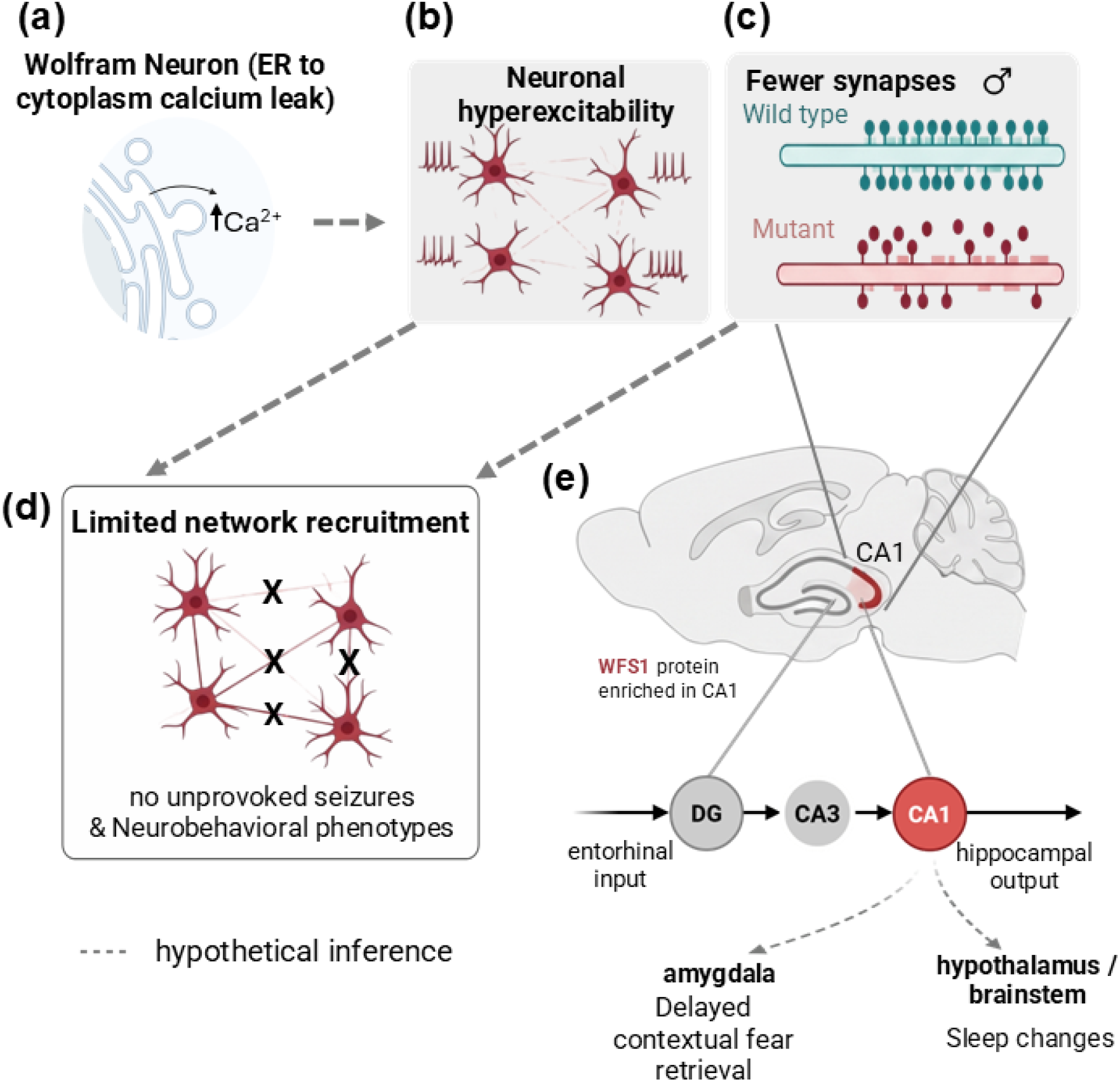
Proposed model of hippocampal network dysfunction in WS mice. Schematic figure summarizing a proposed mechanism linking synaptic, cellular, and network-level findings to their *in vivo* electrophysiological and behavioral phenotypes in WS mice homozygous for *Wfs1*: p.W542*. (a) Loss of wolframin has been reported to cause Ca²□leak from the endoplasmic reticulum (ER) into the cytoplasm. (b) Dissociated hippocampal cultures from male mutant mice show increased neuronal fi ing without a corresponding increase in network synchrony, consistent with a hyperexcitability that arises at the level of individual neurons. We hypothesize that ER Ca² leak contributes to this cellular hyperexcitability. (c) In WS male mutant CA1 stratum radiatum, there is a predominant reduction in excitatory synaptic apposition. (d) We propose that these two features (b, c) act together wherein hyperexcitable neurons embedded in a network with reduced excitatory connectivity generate excitation that is not recruited into coordinated activity, accounting for the absence of unprovoked seizures in mutant mice and in patients. (e) Wolframin is enriched in hippocampal CA1 (red), positioning the CA3 to CA1 pathway as a predicted site of vulnerability. The same reduction in effective connectivity is hypothesized to degrade the organization of hippocampal output, consistent with reduced low-gamma power in male mutants *in vivo*. Reduced connectivity is therefore hypothesized to underlie both the seizure-free phenotype and the impairment of behavior in form of impaired fear memory and sleep changes. Dashed arrows denote hypothetical inferences that were not directly tested.

## Discussion

The neurological phenotype of Wolfram syndrome (WS) has primarily been attributed to dysfunction of the cerebellum and brainstem, the regions where atrophy is most consistently observed on neuroimaging^20,51–53^, even though the hippocampus is among the regions of highest wolframin expression^22,23^, a mismatch we confirm here, with prominent hippocampal CA1 immunoreactivity and low cerebellar signal (figure 1). In WS male mice which are more severely affected, hippocampal wolframin loss causes a predominant reduction of excitatory synapses, while hippocampal neurons become hyperexcitable without generating more network bursting. These findings do translate *in vivo* in our mouse model, where we see an absence of seizure on intracranial EEG, in the absence of hippocampal astrogliosis (Supplementary Figure 6). Additionally, on EEG we found a decrease in low-gamma band in males which co-occurs with REM-exit instability seen in male mice. Together these findings indicate that increased firing at the level of individual neurons does not translate into increased activity at the level of the network.

Loss of wolframin produces two consequences which complementarily act at different levels of neural organization and together shape the network phenotype we observe. First, it reduces excitatory synaptic connectivity wherein we find a predominant loss of excitatory synapses, consistent with reports that *WFS1*-deficient neurons form fewer synapses, extend shorter neurites, and show reduced spontaneous excitatory synaptic transmission^54^. Notably, this loss is likely structural rather than a change in post-synaptic receptor content, as hippocampal GluA1, GluN1 expression were unchanged (Supplementary Figure 3). Second, since wolframin is an ER protein, loss of wolframin leads to cellular calcium dyshomeostasis, and elevation of cytosolic calcium^17,55^, which when paired with *WFS1*-deficient astrocytes releasing excess glutamate through reduced EAAT2 expression^54^, can potentially raise excitatory tone and increase neuronal firing. The relationship between these two effects and network output is context-dependent. For example, in channelopathies like KCNQ2-related disorders, increased excitability at the cellular level^1,2^ does lead to seizures in almost all patients^3^, however cellular hyperexcitability in neurodegenerative conditions like Alzheimer’s disease (AD) does not always lead to seizures, with seizures occurring in about 15% patients^6,7^. Notably, hippocampal synapse loss is a prominent feature in AD^4,5^, raising the possibility that reduced connectivity may limit the translation of cellular hyperexcitability into network-level hyperexcitability. Ultimately, which outcome translates to *in vivo* in mice and then humans is dependent on many factors, and at each level converges on to the level of connectivity substrate that is preserved. Hence, we tested our hypothesis both *in vitro* and *in vivo*, and our data falls in the later class, wherein unit firing is markedly elevated, primarily driven by regular-spiking units, yet network bursting and the network’s distance from criticality are unchanged, and the animals are seizure-free *in vivo*, concordant with the clinical observation that seizures are uncommon in WS patients^9^. Importantly, because this hyperexcitable state is present in dissociated cultures, without any brainstem and cerebellar input, it may represent hippocampal dysfunction at the cellular level and less likely from a remote consequence of degeneration elsewhere.

Hippocampal gamma oscillations depend on fast GABAergic synaptic inhibition, whose decay kinetics together with the level of excitatory drive onto inhibitory neurons set the oscillation frequency, and such rhythms can arise within interneuron networks or through reciprocal pyramidal–interneuron loops^56,57^. Gamma power therefore depends not only on the magnitude of excitatory activity but also on the temporal precision with which excitatory synapses recruit inhibitory neurons into a common rhythm. In our WS male mice homozygous for p.W542*, relative low-gamma power was reduced (Figure 4b). This is interesting because low gamma in CA1 is coherent with CA3 and thought to reflect input from the recurrent network^58^, whose output onto CA1 comprises the Schaffer collateral synapses in stratum radiatum (SR). Our SEQUIN data in males showed the largest loss of excitatory synapses in the SR compartment (Figure 2). Consistent with this link, female mutants, in whom excitatory synaptic loci were not detectably reduced (Supplementary Figure 4), showed no change in low-gamma power (Figure 4c). Because relative power is also influenced by the overall slope of the power spectrum, the male reduction may reflect reduced gamma-band synchrony, a change in the aperiodic background, or both. Hippocampal gamma is also prominent during REM sleep^49,59,60^, and reduced gamma at REM transitions has been reported alongside REM fragmentation in a rodent model of stress vulnerability^61^. In our model, this association is correlational, and our data do not establish that reduced gamma activity causes REM instability. Further work would be needed to specifically define how synaptic connectivity constrains gamma generation in this model. Nonetheless, these findings open the possibility of the use of quantitative non-ictal EEG for discovery of novel biomarkers of network dysfunction in WS patients.

Our findings extend literature on *Wfs1*-deficient mice that has not previously been connected to circuit physiology. The enrichment of wolframin protein in the hippocampal CA1 region has been known for two decades^23,62^, but the circuit function has not been measured. Along the same lines, molecular changes in hippocampus have been studied using transcriptomic profiling of *Wfs1*-deficient hippocampi, which identified several hundred differentially expressed genes, which was an order of magnitude more than the hypothalamus in the same animals^63^, indicating hippocampal transcriptomic dysregulation, but the downstream effect of that on circuit level function was untested. The excitability and sleep phenotype we describe here also has a cross-species precedent. In *Drosophila*, loss of the *wfs1* homolog reduces sleep and dampens circadian rhythm by altering the excitability of wake-promoting dopamine 2-like receptor (Dop2R)-expressing neurons through ER-mediated calcium homeostasis^64^ thereby independently linking wolframin loss, altered neuronal excitability, and disrupted sleep through the known ER-calcium mechanism, and lending evolutionary support to the sleep disruption we see in our animals.

Our data indicate that the synaptic consequences of wolframin loss differs by sex, with excitatory synapses predominantly affected in males and the inhibitory gephyrin scaffold predominantly affected in females (Figure 2; Supplementary Figure 4). Secondary analyses of the intracranial hippocampal EEG showed a corresponding divergence, wherein the aperiodic exponent shifted towards inhibition-dominance in male mutants and excitation-dominance in female mutants (Supplementary Figure 5), mirroring the synaptic phenotype in each sex. Although the histological and electrophysiological measurements were obtained from independent cohorts, their concordant direction within each sex provides convergent evidence for sex-dependent alterations in hippocampal excitation–inhibition organization following wolframin loss. Complementary to our data, sex-dissociated effects on excitatory versus inhibitory synapses have been studied in the hippocampus. For example, developmental anesthetic exposure alters excitatory synaptic transmission in males and inhibitory transmission in females, assessed with the same marker panel we used here^65^, and maternal immune activation produces sex-specific differences in excitatory and inhibitory synapse densities in the dentate gyrus^66^. More broadly, the same perturbation (of tailshocks) can move hippocampal spine density in opposite directions depending on sex, and hormones^67^. At the signaling level, estrogen directly affects ER calcium handling, unfolded protein response via upregulation of SERCA2A^68^. Because wolframin loss acts by depleting ER calcium and dysregulating the unfolded protein response, an estrogen-dependent difference in ER calcium refilling capacity may alter the downstream consequences of *WFS1* loss in a sex-dependent manner. In the context of WS, sex differences in phenotypes are known, with male WS patients showing earlier disease onset^11^, and *Wfs1*-knockout mice showing more severe systemic phenotypes^69^. Because the effects are sex-specific rather than uniform, sex-pooled analyses risk diluting them, and hence similar to our data, sex-stratified analyses should be performed for the neurological phenotype in both model systems and patient registries.

Cerebello-hippocampal projections support oscillatory coupling^24,72^ and may modulate hippocampal dynamics during REM^73^, especially in context of WS. Future work should focus on further delineating this interaction.

Our model carries a homozygous nonsense variant and therefore represents the severe end of the WS genotypic spectrum, which bears on how these findings may translate to patients. Neurological severity in WS varies considerably and is only partly predicted by genotype, though biallelic truncating variants produce earlier and more severe disease than in-frame variants^11,34^. In primary synaptic and neurodevelopmental disorders this genotypic gradient extends to the synapse itself. For example, SHANK2 variants seen in patients, show reduced dendritic synaptic density relative to controls^74^, and across the SHANK family, variant severity may track a gradient of cognitive impairment^75^. Similarly, in POGZ-related disorders, missense variants produce mild phenotypes whereas truncating variants produce severe disease^76^. If a similar relationship holds in WS, the magnitude of hippocampal excitatory synapse loss reported here could represent an upper limit, with patients carrying hypomorphic or in-frame variants showing attenuated synaptic and network phenotypes. This has direct implications for therapeutic trials. Sodium phenylbutyrate and taurursodiol (PB and TURSO), which targets ER and mitochondrial stress, was recently associated with improved or stabilized pancreatic function, vision, and symptom burden in a phase II trial in WS^77^, and an earlier trial of the ER calcium stabilizer dantrolene showed a β-cell signal without measurable change in neurological function^78^. In both cases, limited neurologic outcomes were assessed clinically which may be insensitive to the synaptic and network-level pathology we describe. The role of PB and TURSO in neurodegeneration and synaptic biology is supported by a phase 2a trial in Alzheimer’s disease, in which treatment reduced the synaptic degeneration biomarker neurogranin alongside tau species^79^. Thus, quantitative EEG measures spanning spectral, aperiodic, and sleep-architecture domains may help provide pharmacodynamic neurologic endpoints capable of detecting target engagement before clinical neurological change becomes measurable.

In conclusion, we identify hippocampal network dysfunction as a potential substrate for the neuropsychiatric features of WS, thereby opening up possibilities for future work to discover patient relevant EEG biomarkers, and considering hippocampus as an area of gene therapy target.

## Methods

### Animals and ethics statement

All experimental protocols were reviewed, approved, and performed in accordance with the relevant guidelines and regulations of the Institutional Animal Care and Use Committee (IACUC) of Washington University in St. Louis and were in compliance with US National Research Council’s Guide for the Care and Use of Laboratory Animals, the US Public Health Service’s Policy on Humane Care and Use of Laboratory Animals, and Guide for the Care and Use of Laboratory Animals.

### Sex as a biological variable

Both sexes were studied for *in vivo* electroencephalography (EEG) and sleep analyses (n = 11 male and n = 8 female mice), and for SEQUIN synaptic quantification (n=13 male, and n=9 female mice) data from male and female mice are reported disaggregated throughout the manuscript. Immunoblotting, GFAP immunohistochemistry, *ex vivo* multielectrode array recordings, and behavioral testing were performed in male mice only.

### Quantitative RT-PCR

Mice were euthanized by CO□ inhalation and immediately animals were transcardially perfused with phosphate buffered saline (PBS). Hippocampus was rapidly dissected in ice cold Hibernate-A medium (ThermoScientific, Cat A1247501) and snap-frozen using a dry ice/ethanol bath with a pre-chilled block. Total RNA was extracted with Qiagen RNAeasy minikit (Cat 74104) according to the manufacturer’s instructions, quantified by NanoDrop, and assessed quality using A260/A280 (>1.9), and A260/A230 (>2). RNA was reverse-transcribed with Applied Biosystems™ High-Capacity cDNA Reverse Transcription Kit (Cat 43-688-14). Quantitative PCR was performed on a Quantstudio 6 Pro using PowerUp SYBR Green (cat: A25742) in 20 µl reactions run in triplicates. Primers for *Wfs1* span the exon 7-8 exon-exon junction upstream of the premature termination codon. Forward primer: CTACGCCAGAAGGTGGTGAA; reverse primer: GCCTTGGAGGCTACGTCAAT. Housekeeping GUSB primers: Forward: GAGGATCAACAGTGCCCATT and reverse: AGCCTCAAAGGGGAGGTG. Expression was normalized to GUSB using the ΔΔCt method. Data are expressed as fold change relative to the wild-type mean.

### Western blotting

Hippocampal tissue from 16-week-old mice was homogenized in ice-cold RIPA buffer (Cell Signaling Technology Cat 9806) with protease and phosphatase inhibitors (Cell Signaling Technology Cat 5872S), incubated on ice for 30 min, and cleared by centrifugation at 15,000 × g for 10 min at 4 °C. Protein concentration was determined by Bradford Assay (ThermoScientific 1856209). Each lane on the gel was 25 µg of protein and separated on 4-20% Tris-glycine gels and transferred to 0.45 mm PVDF membranes. Membranes were blocked in 5% BSA in TBS-T for 1 hour at room temperature and incubated overnight at 4 °C with primary antibodies (Supplementary Table 1), followed by HRP-conjugated secondary antibodies for 1 hour at room temperature. Signal was detected with ECL substrate (Biorad 1705061) with normalization to housekeeping markers (β-actin or β-tubulin, supplementary table 1).

The anti-wolframin antibody used (Abcam ab259362) is raised against residues 2–400 of the protein, N-terminal to the W542 truncation site, and therefore detects both full-length wolframin and any stable truncated product and no band of the size predicted for the truncated protein was detected in mutant tissue. n = 3 mice per genotype.

### Immunohistochemistry

Mice were euthanized by CO□ inhalation. Immediately following that, animals were transcardially perfused with phosphate buffered saline (PBS) containing 0.3% heparin. Brains were fixed in 4% paraformaldehyde at 4 °C overnight and then cryoprotected in 30% sucrose solution. Brain was sectioned at 30 µm on a cryostat. Staining was performed free-floating essentially as previously described^80^. Sections were rinsed three times for 5 minutes each in PBS, and then photobleached overnight in PBS containing 4% glycine, 4% acetamide and 0.05% sodium azide in a 10,000 K light box to reduce autofluorescence. Samples were then blocked for 1 hour in 20% normal goat serum (Vector, S-1000-20) with 0.3% Triton X-100 and 0.02% sodium azide in PBS. Sections were incubated with primary antibodies (Supplementary Table 1) for 2–3 days at room temperature in buffer containing 10% normal goat serum, 0.3% Triton X-100 and 0.02% sodium azide in PBS, rinsed three times for 5 min each in PBS, then incubated overnight at room temperature with fluorophore-conjugated secondary antibodies (Supplementary Table 1), protected from light. Sections were rinsed twice 15 minutes each in PBS containing 0.3% Triton X-100, counterstained with DAPI (Invitrogen, D3571; 1:50,000 in 1× PBS) for 20 min, rinsed two times for 5 minutes each in PBS, and mounted on charged slides. Dried sections were coverslipped with ethanol-cleaned coverglass (Marienfeld, 107242) in Mowiol 4-88/AF-300 (9:1; Electron Microscopy Sciences, 17977-150 and 17977-25) and cured in the dark. Images were acquired on a Zeiss LSM 980 with Airyscan or the Synergy H1 with identical acquisition settings applied across genotypes within each imaging session.

### SEQUIN synaptic quantification

Synaptic density was quantified using SEQUIN (Synaptic Evaluation and QUantification by Imaging Nanostructure) as previously described^37,80^, with modifications noted below. Mice aged 16 weeks were perfused and sectioned. Male and female mice were processed and imaged as separate cohorts. Sections containing hippocampus were immunolabelled for the excitatory pair PSD-95 and synapsin, or the inhibitory pair gephyrin and synapsin (antibodies, catalogue numbers and dilutions in Supplementary Table 1). Super-resolution images were acquired on Zeiss Airyscan LSM 980 with identical acquisition parameters applied to all regions of interest (ROIs). Two regions of interest were sampled per animal: CA1 stratum radiatum (SR) and the dentate gyrus molecular layer (ML). For estimating excitatory synapses, synapsin and PSD-95 puncta were detected in 3D, and synaptic loci were defined as pre-/postsynaptic pairs separated by less than 280 nm in three dimensions. For estimating inhibitory scaffold, total gephyrin puncta averaged across all ROIs per animal is reported as previously described^80,81^. Values from all fields within an animal were averaged, and the animal was the unit of replication. All acquisition and analysis were performed blinded to genotype.

### Multielectrode array (MEA) recording

Hippocampi were dissected from P1 male pups in ice cold Hibernate-A medium (ThermoScientific, Cat A1247501), and tissue dissociation and cell plating were performed as previously done^82,83^. Hippocampi were incubated in a dissociation solution containing 6 mg papain (P4762, Sigma, USA) in 3 mL Hibernate E-Ca for 10 min at 30 °C. After replacement of the dissociation solution with Hibernate EB medium (HEB-Ca, BrainBits, USA), the tissue was mechanically triturated using a fire-polished Pasteur pipette to obtain a single-cell suspension. The suspension was centrifuged at 200 X g for 1 min, and the cell pellet was resuspended in Neurobasal Plus medium (A3582901, Thermofisher, USA) supplemented with B-27 supplement (A3582801, Thermofisher, USA), GlutaMAX (35050061, Thermofisher, USA), and penicillin-streptomycin. Before plating, 60-channel microelectrode arrays (MEAs; Multi Channel Systems) were coated with 0.1% polyethylenimine (PEI, Sigma, USA) in water for 30 min, rinsed with sterile water (3 times), and air-dried under the cell culture hood. The arrays were subsequently treated with laminin (20 µg/mL in culture medium) for 30 min. After removal of excess laminin, 300,000 dissociated cells were plated on each MEA. Cultures were maintained at 37 °C in a humidified air containing 5% CO□. Half of the culture medium was replaced with fresh supplemented Neurobasal Plus medium at days *in vitro* (DIV) 4 and every 2 days thereafter. All electrophysiological recordings were performed at DIV 14.

Spontaneous extracellular activity was recorded from the hippocampal cultures using a 60-channel MEA recording system (Multi Channel Systems). During recording, cultures were maintained at 37 °C and 5% CO□ and were allowed to equilibrate on the recording system for approximately 15 min before data acquisition. Signals were acquired at 10 kHz. Raw voltage traces were digitally high-pass filtered at 600 Hz. Negative-going spikes were detected independently on each electrode using a threshold of 5.5 times the channel-specific background-noise estimate, where noise was estimated as the median absolute signal amplitude divided by 0.6745. Successive detections were required to be separated by at least 3 ms, and threshold crossings were aligned to the local voltage minimum within the following 2 ms. Electrodes with a mean firing rate greater than 0.5 Hz over the analyzed recording interval were classified as active and included in subsequent electrode-level analyses.

### MEA data analysis

Mean firing rate was calculated for each active electrode as the number of detected spikes divided by the analyzed recording duration. The mean electrode firing rate for each MEA was calculated by averaging across its active electrodes.

For population-burst analysis, spike trains from all active electrodes were divided into 50-ms bins. The total spike count in each bin was divided by the bin duration and the number of active electrodes to obtain the population firing rate in spikes per active electrode per second. This channel-wise normalization accounted for differences in the number of active recording electrodes among MEAs and enabled comparison of population activity across cultures. The population firing-rate trace was smoothed using a Gaussian filter with a standard deviation of one bin. A population burst was operationally defined as a sustained elevation of the normalized population firing rate above a recording-specific threshold, calculated as the median plus five times the median absolute deviation of the smoothed trace. Consecutive bins at or above this threshold were identified as candidate population bursts. Events lasting at least 50 ms were retained, and events separated by 100 ms or less were merged. Population-burst rate was calculated as the total number of detected bursts divided by the analyzed recording duration in minutes.

For unit-level waveform analysis, detected spikes were aligned to their negative voltage minimum, and 3-ms waveform segments extending from 1 ms before to 2 ms after each spike peak were extracted separately from each electrode. At each waveform time point, amplitude values were standardized across detected spikes by subtracting their mean and dividing by their standard deviation. This placed the waveform features on a comparable scale before dimensionality reduction. The standardized waveforms were projected onto the first two principal components, and a four-component Gaussian-mixture model was applied to the principal-component scores from each electrode to identify candidate unit clusters^84^. Candidate clusters containing 500 or fewer spikes over the approximately 1,000-s recording period, corresponding to a mean firing rate of approximately 0.5 Hz or less, were excluded.

A mean extracellular waveform was calculated for each retained unit and interpolated using cubic interpolation. Trough-to-peak duration was measured from the negative voltage trough to the subsequent positive peak. Units with a trough-to-peak duration of 0.32 ms or greater were classified as regular-spiking units (RSUs), whereas units with a duration below 0.32 ms were classified as fast-spiking units (FSUs). The threshold was determined by bimodal distribution of trough-to-peak durations. The use of extracellular waveform shape to distinguish putative excitatory and inhibitory units followed previous practice^85,86^. Because extracellular waveform shape does not directly establish neurotransmitter identity, RSUs and FSUs were interpreted as putative excitatory and inhibitory units, respectively. Unit firing rate was calculated as the total number of spikes assigned to that unit divided by the recording duration. For each MEA, mean firing rates were calculated separately across retained RSUs and FSUs. Class-level mean RSU and FSU waveforms were generated by averaging the interpolated unit mean waveforms within each class.

Two estimators of the branching parameter were computed. For comparability with the in vitro avalanche literature, a conventional single-step branching ratio (BR) was calculated as the mean ratio of the number of active channels in time bin t+1 (descendants) to the number of active channels in bin t (ancestors), including transitions with zero descendants and excluding bins with zero ancestors, for which the ratio is undefined^87,88^. This parameter was computed across 8 bin-widths (1, 2, 5, 10, 25, 50, 75, 100 ms), applied identically across all recordings. However, because the conventional BR is biased by both bin width^42^, and the fraction of network sampled^43^, it is reported in Supplementary Figure 2 as a comparator and not a primary measure. The primary measure was obtained by multistep regression (MR) estimation, which is invariant to spatial subsampling and is therefore appropriate for array recordings that observe a small fraction of the underlying network^43^, implemented in the mrestimator Python toolbox (version 0.1.8)^41^. Population spike counts pooled across all active electrodes were binned at the same 8 bin widths, and continuous baseline recordings were divided into non-overlapping 5-second trials. Lagged regression coefficients were computed using the stationary-mean method over lags from one bin to 500 ms and fitted with the package’s exponential-with-offset model, yielding the branching parameter m = exp(−Δt/τ), where Δt is the bin width. Instead of a single bin width, m is reported across all 8 bin widths for both genotypes (Fig. 3i). For each genotype, the predicted bin-width dependence of the branching parameter, m(Δt) = exp(−Δt/τ), was generated from the fitted autocorrelation time and overlaid on the measured estimates. Agreement between measured and predicted values indicates that the decline in m with increasing bin width reflects a single underlying autocorrelation time rather than a difference in dynamical regime between genotypes.

### Intracranial electrode implantation and EEG acquisition

Mice of both sexes aged 17-19 weeks were anaesthetized with isoflurane and placed in a stereotaxic frame, with body temperature maintained at 37 °C and depth of anesthesia monitored throughout. Burr holes (0.9 mm) were drilled through the skull over the target locations. Stainless-steel screw electrodes were implanted over the frontal and parietal bones, with ground and reference screw electrodes placed over the interparietal bone. Custom 0.127-mm-diameter platinum–iridium wire electrodes, sheathed in fused silica, were stereotactically lowered through burr holes into dorsal hippocampus and cerebellum following the atlas of Paxinos and Franklin^89^. Bilateral electrodes were implanted into the nuchal muscle for electromyography (EMG). Ground and reference screw electrodes were placed over the interparietal bone. Electrodes were secured with dental acrylic and cyanoacrylate and connected to a headmount. Mice received buprenorphine IP 0.1 mg/kg perioperatively and recovered for 3-5 days before recording.

Recordings were performed via a 16-channel headstage (M4016, Intan Tech) connected to a Stim/Recorder Controller (M4200, Intan Tech). Signals were acquired at 30 kHz and decimated to a stored sampling rate of 468 Hz. Six channels were recorded per animal: bilateral hippocampal depth, bilateral cerebellar, and bilateral nuchal EMG. Continuous EEG and EMG was recorded for three consecutive days per animal, approximately 23 h per day, in the home cage under a 12-hour light/dark cycle, with mice freely moving. The final cohort comprised n = 11 male (5 wild-type, 6 p.W542*) and n = 8 female (4 wild-type, 4 p.W542*) mice. We assessed channels using an automated quality-control procedure and excluded those that failed, and recordings were evaluated for signal quality after acquisition. Power spectral density was calculated using Welch’s method within 30-s epochs with 4-s segments and 50% overlap. Band power was integrated over delta (0.5–4 Hz), theta (4–8 Hz), alpha (8– 12 Hz), beta (13–30 Hz), low gamma (30–50 Hz), and high gamma (50–90 Hz). Relative band power was calculated for each epoch as power within the band divided by total power over 0.5–90 Hz, expressed as a percentage. We averaged values absolute power in decibels and relative power as percentage across quality-passing channels within a recording, then across epochs within each sleep state and light phase, then across recording days. Spectrograms (Figs 4a, 5a) display power in decibels relative to the median of each recording. We compared low-gamma power between genotypes separately within each sex and recording site.

Sleep states were scored in 1-second epochs as wake, NREM, or REM sleep based on spectral features (delta and theta power) together with nuchal EMG amplitude, using Somnotate^48^. Agreement with manual scoring in this cohort was Cohen’s kappa 0.871. We computed sleep architecture measures per recording day, averaged within an animal, and compared them between genotypes separately within each sex. Trained reviewers screened recordings for electrographic seizures by examining the entire record as consecutive two-hour multichannel spectrograms. Isolated interictal epileptiform discharges were not appreciated on review of the record. Quantitative spectral review showed no background changes indicative of seizure activity in animals from either genotype.

Aperiodic (1/f) components were calculated using power spectra which were parameterized into periodic and aperiodic components using specparam/FOOOF algorithm^90^. Welch spectra computed as described above from hippocampal recordings were fitted over 1–45 Hz with settings peak_width_limits = (1,8), max_n_peaks=6, min_peak_height=0.1, peak_threshold=2 and aperiodic_mode was fixed for the reported values (the knee parameter was not fitted, as inspection of log–log spectra showed no knee within the fitting range). The upper limit of the fitting range was set below the 60 Hz notch to avoid contaminating the aperiodic fit. Fits with R² < 0.9 were excluded. The aperiodic exponent α was extracted per epoch and averaged within an animal, with each animal as the unit of replication. The exponent is reported as a positive value, such that a larger α corresponds to a steeper spectral slope and, by convention, an inhibition-dominant shift in excitation/inhibition balance. As above analyses was performed on all tested mice n=11 males and n=8 females (Supplementary Figure 5).

### Behavior Testing

All behavioral testing was conducted in the IDDRC Animal Behavior Subunit at WashU Medicine. For behavioral analysis, separate cohorts of male mice were used to assess phenotypes. Experimenters were blinded to experimental group designations during behavioral testing. All tasks were run by male experimenters during the light phase. Testing orders were randomly counterbalanced for group across apparatuses and trials. Animals were allowed to acclimate to the testing room for 30-60 minutes prior to testing. Unless otherwise indicated, all equipment was cleaned between animals with a 0.02% chlorhexidine diacetate solution (Nolvasan, Zoetis).

Fear conditioning was performed as we previously published^91,92^ and as delineated in figure 6a. Briefly, to condition the animals to associate the context and cue with the aversive foot shock on day 1, the animals were exposed to a 1.0 mA current paired with an 80 dB white noise tone three times on day 1. Contextual fear conditioning was assessed on day 2 by placing the animal back into the chambers from day 1 without shock or tone presentation. On day 3, the animals were placed into a contextually altered chamber (dimensions, odorant, materials) and assessed for cued fear conditioning in the presence of the 80 dB tone from day 1. On all three days, freezing was quantified by FreezeFrame software (Actimetrics). Sensitivity to foot shocks was evaluated one week following the start of conditioned fear testing to confirm differences in freezing behavior were not confounded by differences in reactivity to the shock current, based on our previously published methods^92^. Mice were placed in the fear conditioning chamber and exposed to a 3 sec foot shock every 20-30 sec starting at 0.05 mA and increasing in intensity by 0.05 mA until a flinch was observed. The current at which this behavior occurred was recorded.

Normal and complex activity wheel was performed based on prior methods^93^. The apparatus is as previously described^93^, including a suspended running wheel, a thin layer of bedding, and a sensor to detect wheel rotations. This test consisted of two segments: normal running wheel and complex running wheel. The normal activity wheel assessed the animal’s ability to traverse a fully intact wheel. The complex running wheel provided a more sensitive measure of fine motor coordination between the fore- and hindlimbs by removing rungs to create irregularly spaced intervals between rungs. Each test segment consisted of a 1-hour test daily for five consecutive days. The test segments were spaced three days apart. The total distance traveled on the wheel, max speed achieved, and total wheel rotations were quantified.

Accelerating rotarod was done based on previous methods^94,95^. The AccuRotor EzRod apparatus (Omnitech Electronics, Inc) was used. Animals received three trials a day on each of four consecutive days on an accelerating rod (4-40 rpms) for 300 seconds. An intertrial interval of 30 minutes was used each day. Each animal received a habituation trial on a continuously rotating rod at 3rpms for 60 seconds prior to the first test trial on the first day only. The latency the animal could remain on the rod, distance traveled and speed achieved was measured by Fusion software (Omnitech Electronics, Inc).

### Statistics

The animal or independently plated cultures were the unit of replication throughout. Units, puncta, ROIs, and events were averaged within an animal or culture before statistical comparison. Exact n values for every panel are given in the figure.

Given the non-normal distributions of several measures, genotype comparisons used two-sided Mann– Whitney U tests. Analyses were performed in SPSS (for behavioral data) and GraphPad Prism v11 for all other datasets. Data are presented as median with individual animal values overlaid unless stated otherwise. P < 0.05 was considered significant and exact P values reported throughout except where P < 0.001.

## Funding

S.A. was funded by KL2 career development award KL2TR002346; UpLifting Athletes Young investigator award; and has clinical trial support from Amylyx Pharmaceuticals Inc., which is unrelated to this work. FU previously had a sponsored research agreement with and received material support from Prilenia Therapeutics; serves as PI of the HELIOS clinical trial supported by Amylyx Pharmaceuticals; has received grants from the National Institutes of Health (NIH; DK132090, DK020579, and HD103525); has received royalties from Novus Biologicals and Sana Biotechnology; has received licensing and/or consulting fees from Opris Biotechnologies and Emerald Biotherapeutics; has received travel support from Wolfram France, Wolfram UK, and the Snow Foundation; serves in unpaid advisory roles for the Snow Foundation and Be A Tiger Foundation; and holds U.S. patents 9,891,231, 10,441,574, and 10,695,324. S.M. and M.W. were supported by Eunice Kennedy Shriver National Institute Of Child Health & Human Development of the National Institutes of Health grant P50 HD103525 to the Intellectual and Developmental Disabilities Research Center at Washington University; National Institutes of Health grant P30 DK020579. AD and SS are supported by the Air Force Office of Scientific Research (FA95502310461) and the National Science Foundation (CBET-2331330). None of the other authors have any conflicts of interest related to this work.

## Acknowledgements

We are thankful to all patients and families with Wolfram Syndrome. We are also thankful to all members of Dr. Urano’s lab, especially Cris Brown.

## Supplementary Figures

**Supplementary figure 1:**
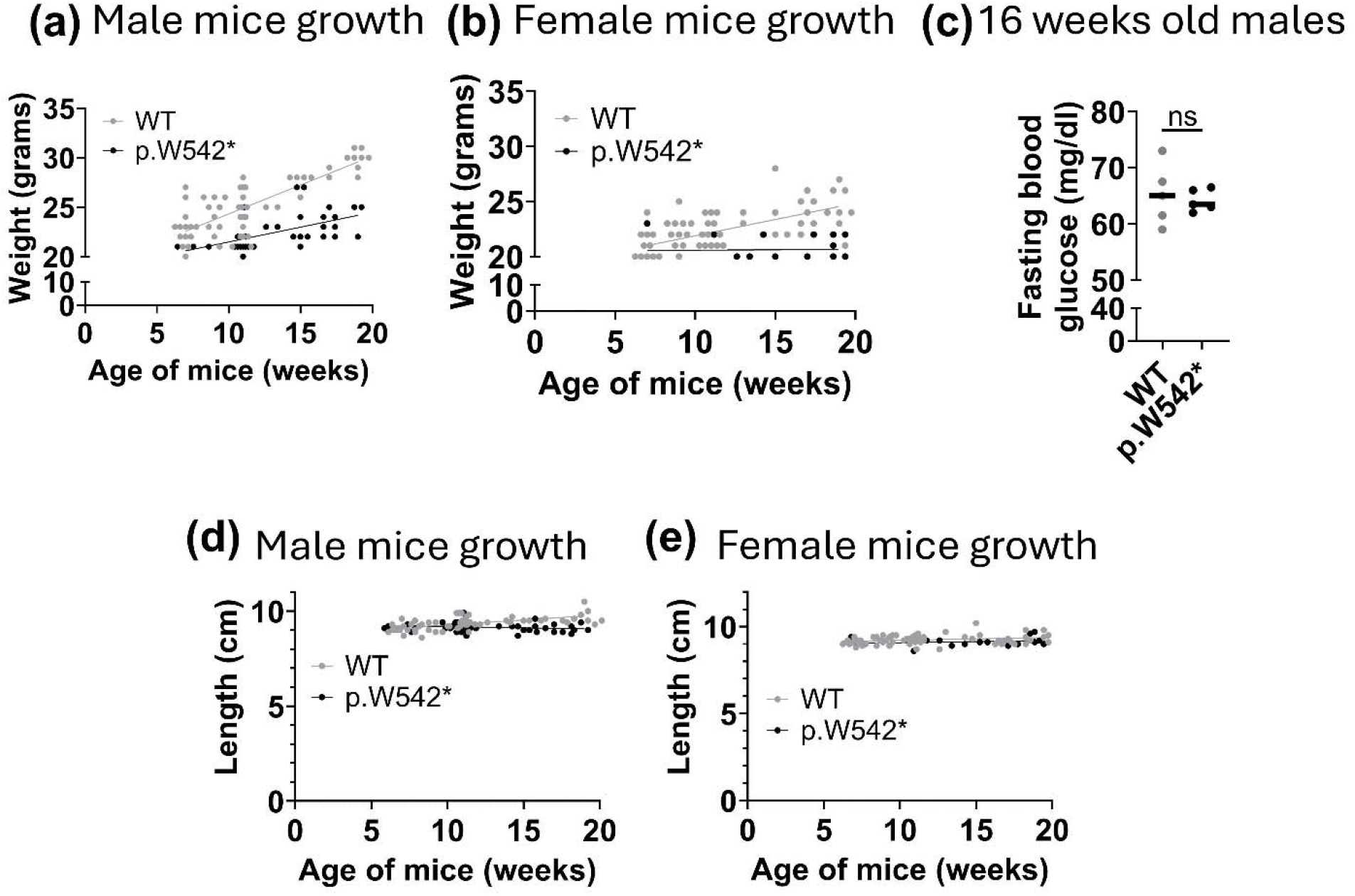
Reduced body weight gain in *Wfs1* p.W542 mice without fasting hyperglycemia. **(a, b)** Body weight from 6-20 weeks of age in male and female mice, with grey lines representing wild-type (WT) and black lines representing *Wfs1* p.W542* homozygous mice. Each data point represents a single animal. Males (WT n = 78, p.W542* = 38), females (WT n = 86, p.W542* = 29). Genotype effects were estimated by linear regression with age as a covariate and standard errors clustered on dam. Homozygous *Wfs1* p.W542* mice were lighter than WT in both sexes. (a) Males weight difference of −3.65 g (95% CI −4.91 to −2.39; p = 1.3 × 10□□) and (b) females, weight difference of −4.07 g (−5.15 to −2.98; p = 2.1 × 10□¹³). **(c)** Fasting blood glucose in 16-week-old males measured using a glucometer. Each data point is a different animal. Mann-Whitney test with p > 0.99. **(d-e)** Naso-anal length of mice against age (6-20 weeks), with each data point representing a different animal. Genotype effects were estimated by linear regression with age as a covariate and standard errors clustered on dam. Homozygous *Wfs1* p.W542* mice were shorter than WT in both sexes. (d) males length −0.38 cm (−0.53 to −0.23; p = 7.7 × 10□□) (e) females length −0.39 cm (−0.54 to −0.24; p = 3.1 × 10□□).

**Supplementary figure 2:**
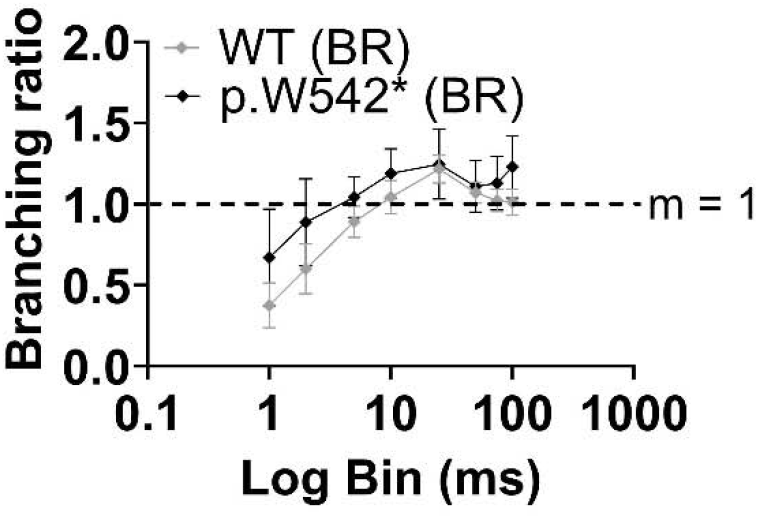
The conventional single-step branching ratio estimator is bin-width dependent and does not resolve a genotype difference. Branching ratio (BR) estimated by the conventional single-step method as a function of analysis bin width (1–100 ms, log scale) for wild-type (grey, n = 4) and Wfs1 p.W542* homozygous (black, n = 4) male hippocampal cultures at DIV14. Datapoints represent mean +/-SD. BR estimates rise monotonically with bin width in both genotypes and cross m = 1 at intermediate bins (20–50 ms), a known consequence of single step branching estimates^42^, and spatial subsampling^43^, rather than a change in the underlying network state.

**Supplementary Figure 3:**
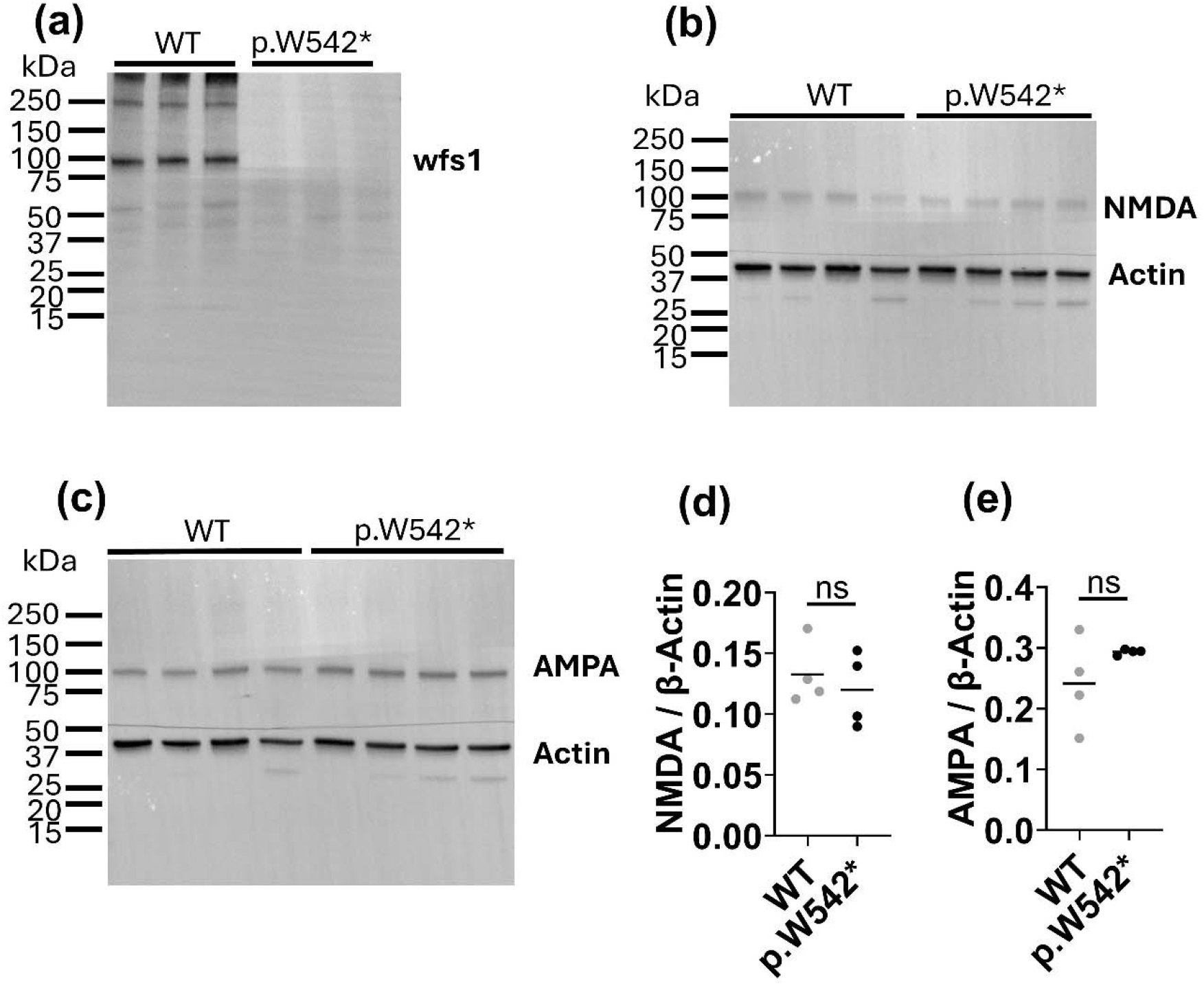
Hippocampal protein expression of WFS1, NMDA and AMPA receptor expression in WT and *Wfs1* p.W542* homozygous male mice. **(a-c)** Representative full western blots showing (a) wfs1 (cropped image shown in figure 1) (b) NMDA receptor 1 (GluN1) (c) AMPA receptor 1 (GluA1). Each sample lane represents a separate male animal hippocampus. β-Actin was used as loading control. Molecular mass markers are indicated in kilodaltons (kDa). **(d, e)** Densitometric quantification of (d) NMDA receptor and (e) AMPA receptor band intensities, normalized to the corresponding β-actin signal. Gray symbols indicate WT samples, and black symbols indicate p.W542* homozygous male samples. Each point represents a separate male animal. Horizontal lines indicate mean. Statistical analyses was performed using Mann-Whitney test, and ns represents non-significant change with panel (d) p = 0.68, and panel (e) p = 0.34.

**Supplementary Figure 4:**
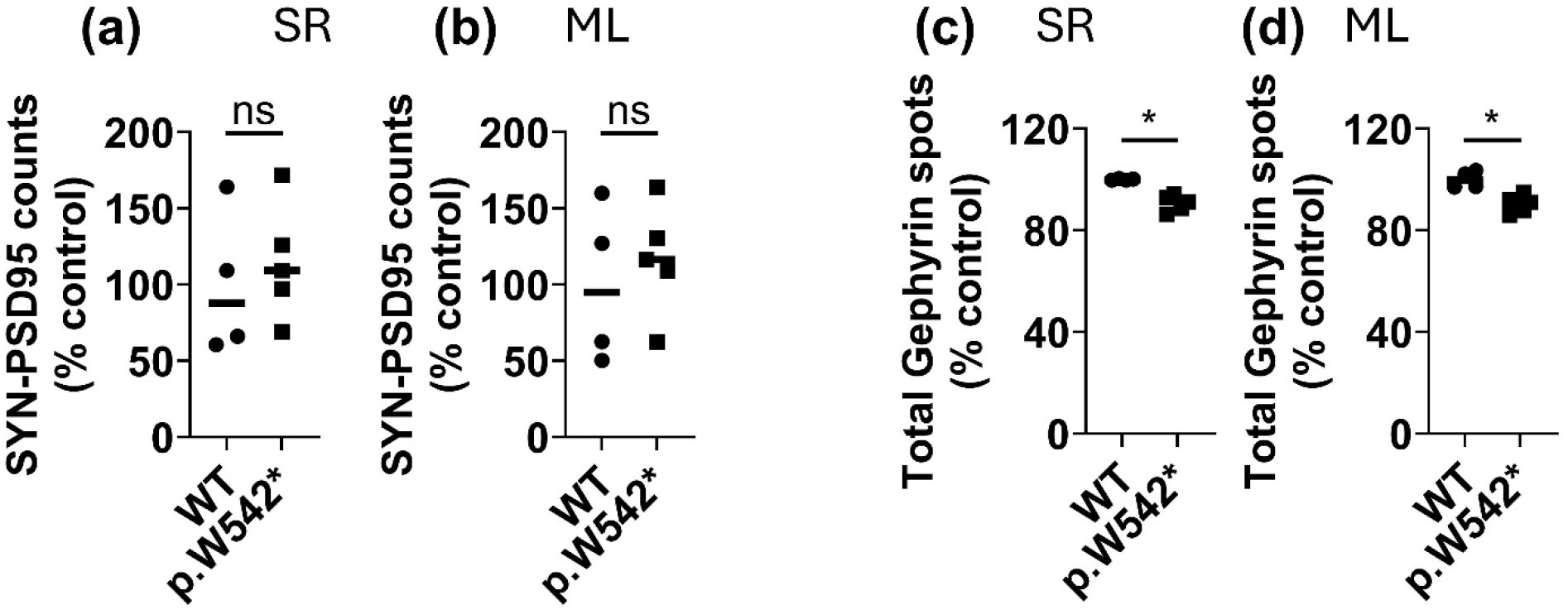
Hippocampal *Wfs1* W542* reduces total gephyrin puncta in females, without a detectable reduction in excitatory synaptic loci. **(a, b)** Female mice, SEQUIN quantification of colocalized synapsin–PSD-95 pairs (excitatory synaptic loci) per region of interest. Values are normalized to the wild-type group mean. (a) SR of CA1, ns: p=0.56; difference in median =+21%. (b) ML of the dentate gyrus, ns: p=0.73; difference in median =+21%. (c, d) Female mice, total gephyrin puncta per region of interest in SR and ML. Values are normalized to the wild-type group mean. Each symbol represents one animal, with each data point representing a mean across all ROIs imaged in that region for that animal (12–24 ROIs per animal for SR, 6-7 ROIs per animal for ML); horizontal bars indicate group medians. n=4 wild-type (WT) female mice and n=5 Wolfram homozygous (p.W542*) female mice. Statistical comparisons by Mann-Whitney test: (c) SR, p=0.016; difference in median = -8.7%. (d) ML, p=0.016; difference in median = -8.8%.

**Supplementary Figure 5:**
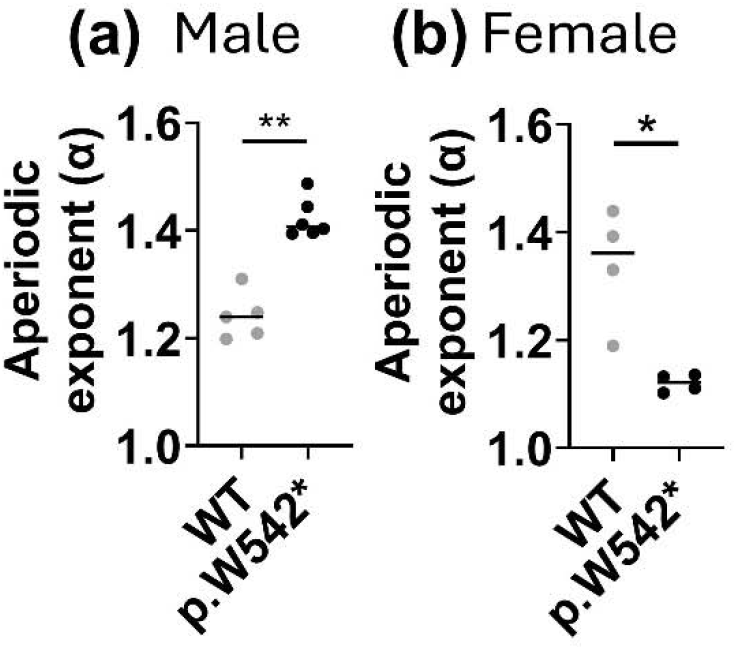
Increased hippocampal aperiodic exponent in male and decreased exponent in female *Wfs1* W542* mice. **(a, b)** Aperiodic exponent of the hippocampal power spectrum, fit by specparam (FOOOF – Fitting oscillations and one-over f) larger α indicating a steeper spectrum and an inhibition-dominant shift^96^. Statistical comparisons by Mann-Whitney test, (a) male **p=0.0043, difference in medians = 0.16 (WT: 1.24; p.W542*: 1.4) (b) female *p=0.028, difference in medians = -0.24 (WT: 1.36; p.W542*: 1.12). The genotype effect differed between sexes (genotype × sex interaction, permutation p < 0.001).

**Supplementary Figure 6:**
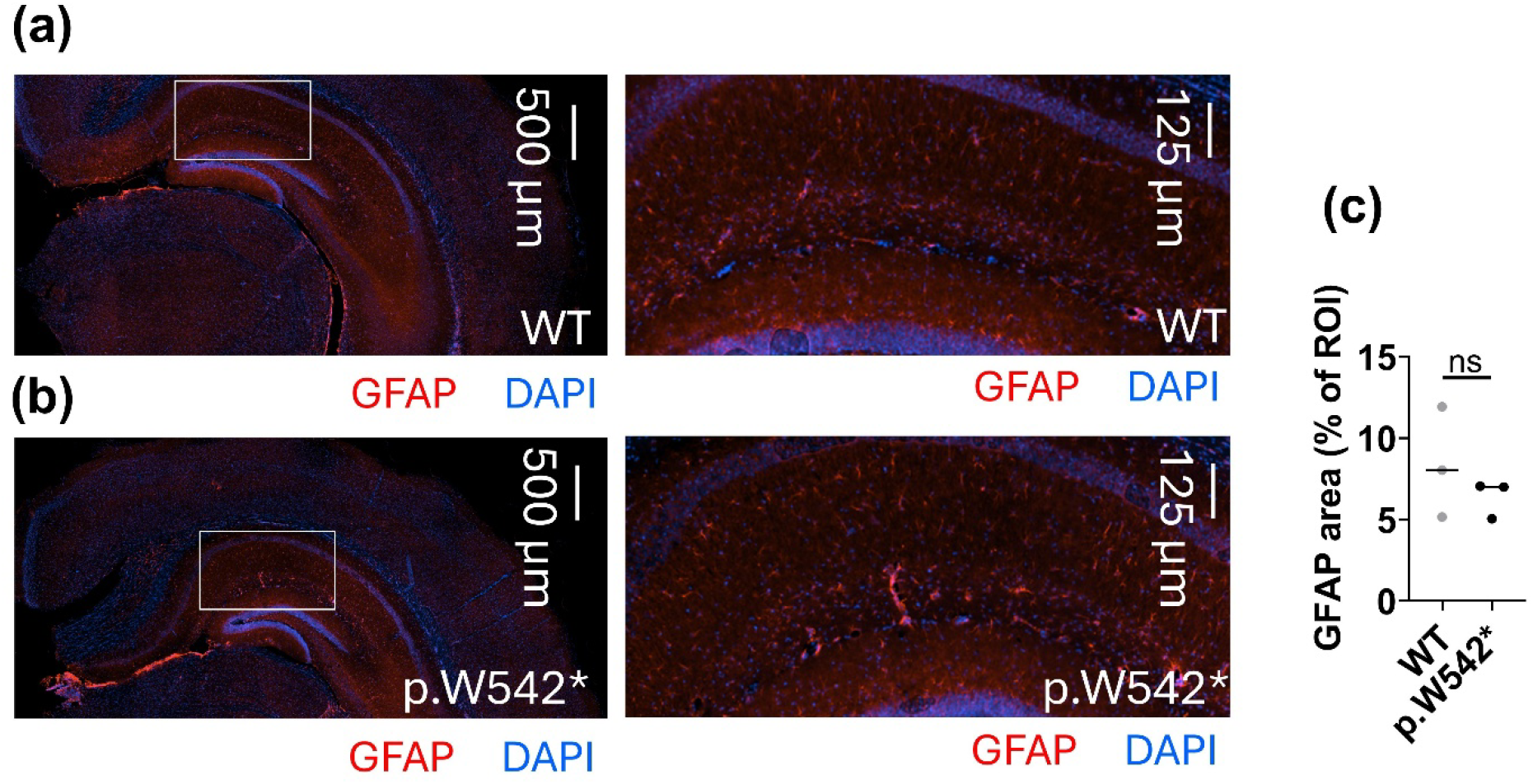
Hippocampal GFAP reactivity unchanged in *Wfs1* W542* homozygous mice. **(a, b)** Representative sections from (a) wild-type and (b) *Wfs1* p.W542* male mice at 18 weeks, immunostained for GFAP (red) with DAPI (blue). Left panel is a hippocampus overview with white rectangle indicating the field enlarged on the right panel, in the CA1 region of the hippocampus. Images were acquired with identical settings and are displayed with identical brightness and contrast. **(c)** GFAP positive area expressed as a percentage of a fixed region of interest positioned in CA1 stratum radiatum, 390 × 279 µm; 108,774 µm², quantified in Fiji (ImageJ) from one ROI per animal. Each point represents one animal; horizontal line indicates the median. WT 8.03% (range 5.16–11.93); *Wfs1* p.W542* 6.99% (range 5.05–7.04). Exact two-sided Mann-Whitney p = 0.40. n=3 animals per genotype. ns, not significant. The absence of detectable astrogliosis is consistent with the absence of epileptiform activity and overt neurodegeneration in this model but does not exclude subtle astroglial changes.

**Supplementary table 1:** Primary and secondary antibodies used in the manuscript, with host species, clonality, commercial source, catalog number, Research Resource Identifier (RRID), and working dilution. ^a^ Two goat anti–rabbit secondary antibodies were used in separate labeling panels: Alexa Fluor 568 for detection of PSD-95 in the SEQUIN synaptic panel, and Alexa Fluor 594 for detection of WFS1 protein. The two panels were never combined on the same section. ^b^ RRID not yet assigned or not confirmed in the Antibody Registry at the time of submission.

| Antibody | Host, clonality | Source | Catalog no. | RRID | Dilution |
| --- | --- | --- | --- | --- | --- |
| **Primary antibodies** |  |  |  |  |  |
| Anti-synapsin 1/2 | Guinea pig, polyclonal | Synaptic Systems | 106 004 | AB_1106784 | 1:1,000 |
| Anti-PSD-95 | Rabbit, polyclonal | Invitrogen | 51-6900 | AB_2533914 | 1:200 |
| Anti-gephyrin | Chicken, recombinant monoclonal IgY | Synaptic Systems | 147 009 | —<sup>b</sup> | 1:500 |
| Anti-WFS1 (wolframin), clone EPR23801-91<sup>a</sup> | Rabbit, recombinant monoclonal | Abcam | ab259362 | —<sup>b</sup> | 1:1000 |
| Anti-WFS1 (wolframin) | Rabbit, polyclonal | Proteintech | 26995-1-AP | AB_2880717 | 1:250 |
| Anti-NMDA Receptor 1 (GluN1) | Rabbit, monoclonal | Cell Signaling Technology | 5704 | AB_1904067 | 1:1000 |
| Anti-AMPA Receptor 1 (GluA1) | Rabbit, monoclonal | Cell Signaling Technology | 13185 | AB_2732897 | 1:1000 |
| Anti-β-Actin | Mouse, monoclonal | Cell Signaling Technology | 3700 | AB_2242334 | 1:1000 |
| Anti-GFAP | Rabbit, polyclonal | Invitrogen | PA3-16727 | AB_2109795 | 1:500 |
| Anti-α-tubulin | Rabbit, monoclonal | Cell Signaling Technology | 2128 | AB_823664 | 1:1000 |
| **Secondary antibodies** |  |  |  |  |  |
| Goat anti–guinea pig IgG (H+L), Alexa Fluor 488 | Goat, polyclonal | Invitrogen | A-11073 | AB_2534117 | 1:200 |
| Goat anti–rabbit IgG (H+L), Alexa Fluor 568<sup>a</sup> | Goat, polyclonal | Invitrogen | A-11011 | AB_143157 | 1:200 |
| Goat anti–chicken IgY (H+L), Alexa Fluor Plus 647 | Goat, polyclonal | Invitrogen | A32933 | —<sup>b</sup> | 1:200 |
| Goat anti–rabbit IgG (H+L), Alexa Fluor 594<sup>a</sup> | Goat, polyclonal | Invitrogen | A-11012 | AB_2534079 | 1:200 |

